# The Specious Art of Single-Cell Genomics

**DOI:** 10.1101/2021.08.25.457696

**Authors:** Tara Chari, Lior Pachter

## Abstract

Dimensionality reduction is standard practice for filtering noise and identifying relevant features in large-scale data analyses. In biology, single-cell genomics studies typically begin with reduction to two or three dimensions to produce ‘all-in-one’ visuals of the data that are amenable to the human eye, and these are subsequently used for qualitative and quantitative exploratory analysis. However, there is little theoretical support for this practice, and we show that extreme dimension reduction, from hundreds or thousands of dimensions to two, inevitably induces significant distortion of high-dimensional datasets. We therefore examine the practical implications of low-dimensional embedding of single-cell data, and find that extensive distortions and inconsistent practices make such embeddings counter-productive for exploratory, biological analyses. In lieu of this, we discuss alternative approaches for conducting targeted embedding and feature exploration, to enable hypothesis-driven biological discovery.

## Introduction

The high-dimensionality of “big data” genomics datasets has led to the ubiquitous application of dimensionality reduction to filter noise, enable tractable computation, and to facilitate exploratory data analysis (EDA). Ostensibly, the goal of this reduction is to preserve and extract local and/or global structures from the data for biological inference [1–3]. Trial and error application of common techniques has resulted in a currently popular workflow combining initial dimensionality reduction to a few dozen dimensions, often using principal component analysis (PCA), with further non-linear reduction to two dimensions using t-SNE [4] or UMAP [1, 2, 5, 6]. For single-cell genomics in particular, these embeddings are used extensively in qualitative and quantitative EDA tasks which fall into four main categories of applications (Table 1, ‘Application’):

- Modality-Mixing, Integration, & Reference Mapping: Embeddings are used to visually assess the extent of integration, mixing, or similarities between cells from different batches [7–9] and to compare methods of integration/batch-correction [10]. For query dataset(s) mapped onto reference datasets/embeddings, visuals likewise provide an assessment of merged data similarities or differences [11, 12].
- Cluster Validation & Relationships: Visual applications range from assessing the existence of and relationships between predefined clusters, to inferring properties of the clusters (e.g. spread or heterogeneity) [1, 2, 13], and to generating the clusters themselves from the two-dimensional space (e.g. to define cell types or detect doublets) [3, 14, 15].
- Density-based Visuals & Marker Analysis: Embeddings are used to justify or measure changes in cell populations between different conditions, by comparing contour locations and sizes in the density diagrams, as well as changes in intensity or spread of gene expression [16–20].
- Trajectory Inference & Continuous Relationships: Embedding applications range from implying or inferring local, continuous relationships between cells and assigning pseudotime coordinates [21–24], to using the two-dimensional coordinates for explicit calculations of magnitude and direction of developmental progression [23, 25, 26].

**Table 1:**
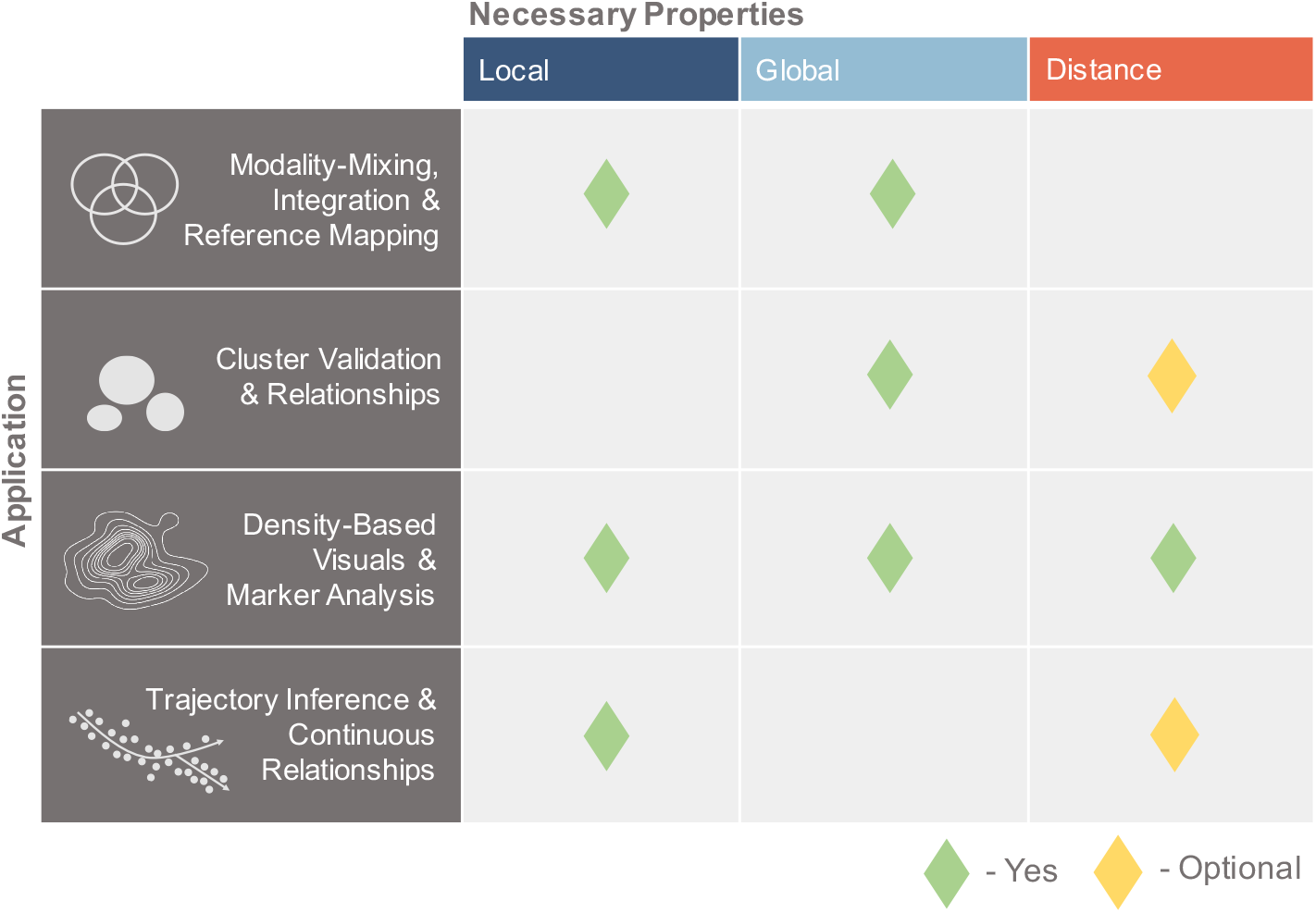
Necessary Properties for Embedding Applications. Application rows denote biological tasks, and columns denote which properties are necessary to preserve for each task.

Inherent in these applications are assumptions of preservation of local and global cell properties, as well as distances, delineated in Table 1. Based on previous works [6, 13, 27, 28] and the objective functions of UMAP and t-SNE [4, 5], ‘local’ is defined as nearest neighbor relationships, ‘global’ as neighbor relationships and properties of groups of cells (e.g. cell types), and ‘distance’ as Euclidean distance (*L*_2_ norm) or Manhattan distance (*L*_1_ norm) between points. Note that preservation of distance implies preservation of local and global properties. We utilize the *L*_2_ norm as it is the default metric of UMAP/t-SNE. We also present results with the *L*_1_ norm (shown in Supplementary Materials), as *L*_1_ is more suitable for measuring distance in high dimensions, particularly in comparison to other L norms [29, 30], and is commonly applied to transcriptomic data [31–33], with comparable performance to the probabilistic Jensen-Shannon divergence in single-cell distance calculations [34].

Yet, despite the goals of these methods [2, 3, 6] to preserve local and/or global structure, there is little theory or empirical analysis to support these claims. For example, while the popular t-SNE and UMAP methods claim faithful representation of local and/or global structure in low dimensions [1, 2, 5], there is evidence they fail in this regard [1, 35], and theorems providing guarantees on the embeddings rely on numerous assumptions unlikely to hold in practice and ignore the coupling of PCA to non-linear methods [36].

Here we assess dimensionality reduction for single-cell gene expression, first investigating the preservation of the necessary properties comprising the columns of Table 1, then assessing the impact of these embeddings across the applications comprising the rows of Table 1.

## Preservation of Local and Global Structure in 2D Embeddings

We begin with the columns of Table 1, and assess the preservation of these properties by twodimensional embedding, as compared to the ambient space or higher-dimensional PCA space to which the 2D reduction is coupled (see Supplementary Methods). ‘Ambient’ space refers to the gene count matrix after highly variable gene selection and log-normalization of the counts (see Supplementary Methods). ‘Coupling’ refers to the combination of a higher-dimensional reduction by PCA followed by a (non-linear) reduction to 2D (e.g. ‘PCA-50D→UMAP’).

### Local Preservation

Given the focus on preserving local nearest neighbors in both the UMAP and t-SNE methods, we first measured the recapitulation of nearest neighbors in 2D embeddings, as compared to the neighbors defined in ambient space. We used Euclidean (L_2_) distance, the default for the non-linear reduction methods, to define each cell’s 30 nearest neighbors and measured Jaccard distance (dissimilarity) between the neighbors in embedding and ambient space (where 1.0 denotes no overlap). We then reduced a (10x Genomics assayed) mouse ventromedial hypothalamus (VMH) neuron dataset [37], an ex-utero cultured mouse embryo dataset (at the E8.5 stage) [8], and a mouse primary motor cortex (MOp) dataset [38] to two dimensions. The 2D t-SNE/UMAP embeddings (e.g. ‘PCA-50D→UMAP’ in Fig. 1a) displayed large Jaccard distances with respect to the neighbors in ambient dimension, with an average consistently above 0.7 (70%) with or without PCA-coupling, and dissimilarity increasing with the size of the dataset (Fig. 1a, Supplementary Fig. 1). The distortion of neighbors was worse for two-dimensional PCA (‘PCA-2D’), consistent with other findings on the poor preservation of local neighborhoods by both PCA and t-SNE/UMAP non-linear reduction methods [1, 35] (top plots, Supplementary Fig. 1). Similarly poor neighbor retention was observed even in the higher dimensional PCA spaces (‘PCA-50D’ Fig. 1a i, Supplementary Fig. 1) [35], and in the reduction from PCA space to the 2D embedding (Fig. 1a ii, right panels Supplementary Fig. 1).

**Figure 1:**
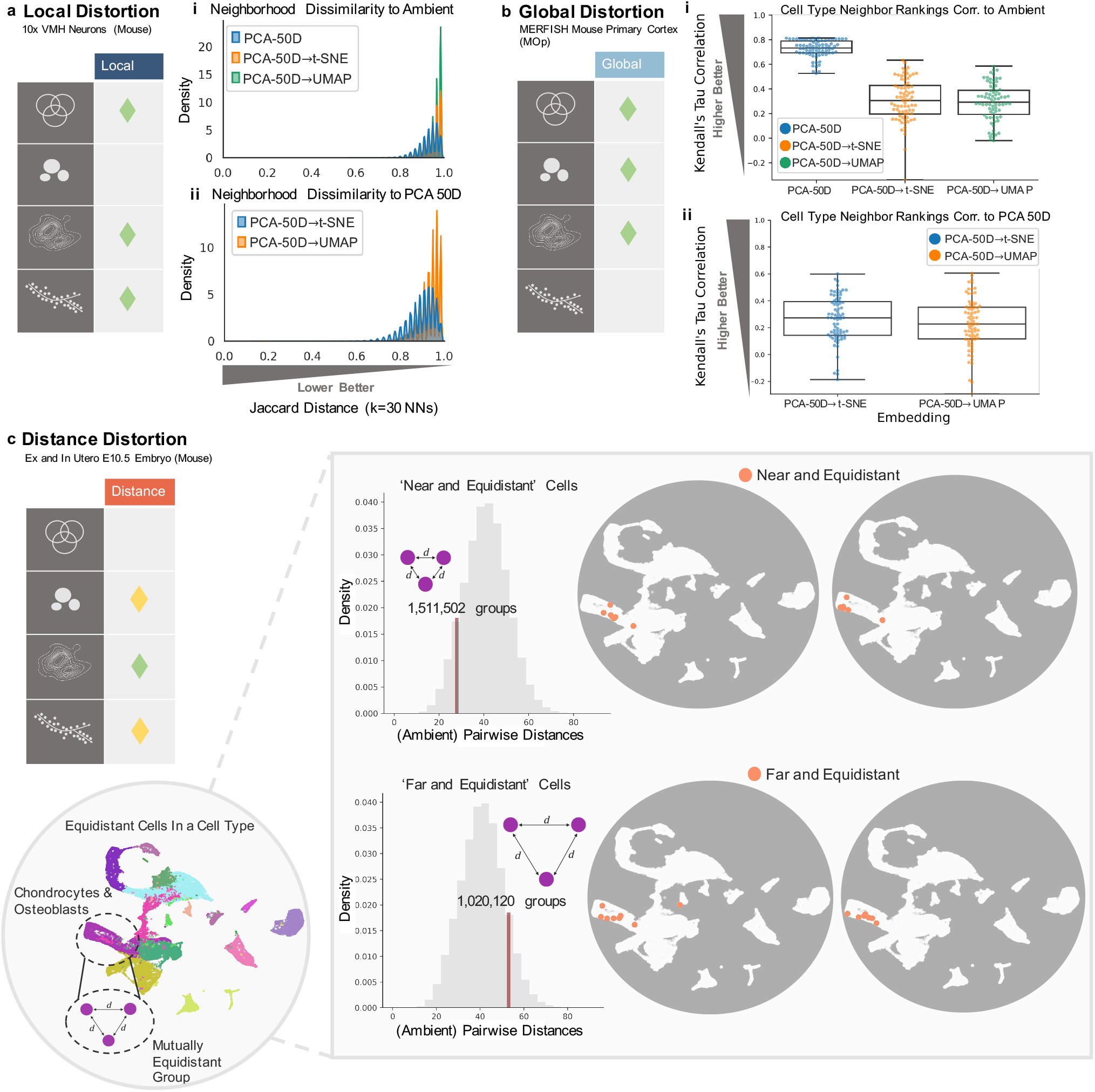
Distortion of Necessary Properties in Embeddings. **a)** i. Distribution of Jaccard distance of cell neighbors in high and low dimensional embeddings, as compared to ambient space. ii. Distribution of Jaccard distance of cell neighbors in low dimensional embeddings, as compared to higher dimensional PCA space. **b)** i. Boxplot of correlations of cell type neighbor rankings to ambient space over three replicate embeddings. ii. Boxplot of correlations of cell type neighbor rankings to higher dimensional PCA space over three replicate embeddings. **c)** Selection of equidistant groups with ‘near’ or ‘far’ distances. UMAP embedding of the data in grey circles, with orange circles denoting all cells within one equidistant group.

### Global Preservation

Turning to global relationships, we measured the preservation of the rankings of neighbors of cell types rather than individual cells. Rankings were constructed from average pairwise distances between the cells of the different types, across replicate 2D embeddings (see Supplementary Methods). For the same datasets, correlation of cell type neighbor rankings to that of the ambient space were low (≤ 0.4), and at least 33% lower than in the higher dimensional PCA spaces, with warped or even reversed correlations in comparison to the ambient (Fig. 1b i, Supplementary Fig. 2) or coupled PCA space (Fig. 1b ii, Supplementary Fig. 2). These distortions were not specific to the distance measure used; we observed similar results when using the *L*_1_ norm to determine cell type neighbors (Supplementary Fig. 2). This is consistent with observations made in other studies [6, 28]. For analyses of recapitulation of cluster properties such as inferred heterogeneity or spread, see ‘Clustering Validation & Relationships’ and ‘Arbitrary Shapes Preserve Structure’ below.

### Distance Preservation

To examine distance preservation, we extracted groups of cells with quantitatively distinct relationships in the ambient space of the Seurat-integrated [7] ex- and in-utero mouse embryo dataset (at the E10.5 stage) from [8], specifically equidistant groups of cells, where the distances between cells were either small (‘near’) or large (‘far’) (Fig. 1c) (see Supplementary Methods). This revealed upwards of 2.5 million such groups, with 3 to 8 cells in each (Supplementary Fig. 3a,e). However, once embedded into two dimensions, these groups of cells (orange dots on UMAPs, Fig. 1c) display the same dispersion patterns, violating distance preservation, and rendering these distinct, transcriptomic relationships indistinguishable.

This is not surprising, given previous theoretical work on the limits of distance preservation in low dimensions, particularly for equidistant points [39–41]. The Johnson-Lindenstrauss Lemma on the optimality of linear embedding [42–44] shows that preservation of pairwise distances with a margin of error of at most 20% for a modestly sized dataset of 10,000 cells would require at least 1,842 dimensions [45]. Distortion is inevitable: given *n* points embedded in two dimensions, the distortion of the ratio of the maximum distance, *D*, to minimum distance, *d* (‘max/min ratio’), grows as 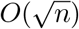 [46] (Supplementary Note, Supplementary Methods).

In practice, measuring these ‘max/min ratios’ in 2D embeddings, for the ex- and in-utero data as well as the 10x VMH neurons, revealed 4- to 200-fold increases in these ratios whether compared to the upstream PCA space or ambient space (with or without PCA-coupling), within groups of equidistant cells as well as nearest neighbors (Supplementary Fig. 3,4). Higher dimensional PCA spaces largely maintained similar max/min ratios to the ambient space (Supplementary Fig. 4). However, we note that in low dimensions PCA embedding of equidistant points is tantamount to applying a random projection, resulting in projected points displaying numerous mirages of structure or outliers (Supplementary Fig. 5).

### Distortion of Trends in Applications

Given the distortions of the necessary properties in Table 1, we then investigated their impact on each row or application i.e. how in practice such embeddings affect the inferences and implications made in each application.

### Modality-Mixing, Integration, & Reference Mapping

Malleability of ‘structure’ under low dimensional embedding is particularly apparent in the mixing properties of integrated, mapped, or batch-corrected datasets, where an integration procedure is accompanied by an embedding of the melded datasets (Fig. 2, Supplementary Fig. 6) [7, 8]. This relies on preserving both local relationships i.e. which cells are mixed, and global patterns i.e. overall trends of mixing or non-mixing between datasets (Table 1). For the integrated ex- and in-utero dataset (E10.5), we calculated the fraction of each cell’s nearest neighbors with the same label as the cell, to compare whether embeddings accurately reflect the extent of mixing of ex- and in-utero cells by integration (Fig. 2a) (see Supplementary Methods).

**Figure 2:**
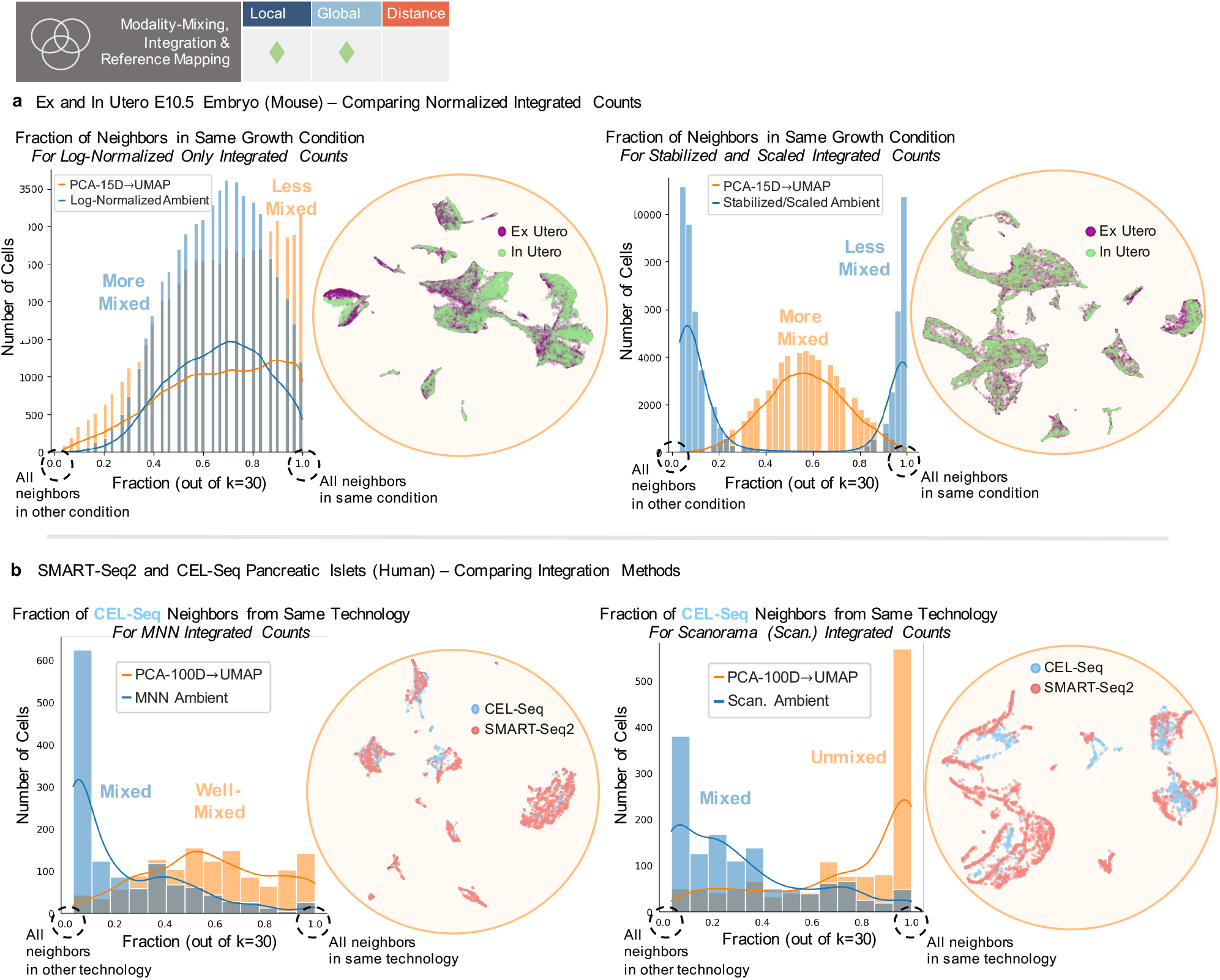
Distortion of Mixing Patterns. **a)** Left plot shows ‘Log-normalized’ ambient (blue) and 2D embedding (orange) distributions of mixing (fraction of cell neighbors in the same condition), where 1.0 is no mixing. Corresponding UMAP shown next to it. Right plot shows ‘Variance Stabilized and Scaled’ ambient (blue) and 2D embedding (orange) distributions of mixing (fraction of cell neighbors in the same condition). Corresponding UMAP shown next to it. **b)** Left plot shows ‘MNN Integrated’ ambient (blue) and 2D embedding (orange) distributions of mixing (fraction of cell neighbors in the same condition) for CEL-Seq cells. Corresponding UMAP shown next to it. Right plot shows ‘Scanorama Integrated’ ambient (blue) and 2D embedding (orange) distributions of mixing (fraction of cell neighbors in the same condition) for CEL-Seq cells. Corresponding UMAP shown next to it.

The ‘Log-Normalized’ integrated, ambient data displayed a largely unimodal, well-mixed distribution of cells between conditions, while the distribution generated from embedding into two dimensions was shifted towards unmixed (left side, Fig. 2a). The ‘Variance-stabilized and Scaled’ integrated, ambient data (scaling performed after integration and log-normalization) displayed the opposite trend. The ambient data presented a bimodal distribution with completely unmixed cell populations, while the final embedding displayed a unimodal distribution of well-mixed cells from both conditions (right side, Fig. 2a). These additions or losses of mixing properties by 2D embedding were replicated using the *L*_1_ metric for neighbor determination (Supplementary Fig. 6a).

Such mixing patterns are not only used to argue that different datasets are similar, but also to argue for the superiority of one integration method over another. To assess whether such inferences are legitimate, we merged the SMART-Seq2 and CEL-Seq pancreatic islet datasets utilized in [9, 10] with one of two methods, MNN [47] or Scanorama [10]. Looking at the fraction of mixing of CEL-Seq cells in the merged ambient space reveals similar mixing by both methods (CEL-Seq cells placed near SMART-Seq2 cells) (ambient distributions, Fig. 2b). However the UMAP embeddings provide opposite pictures, with MNN appearing to result in a well-mixed distribution of CEL-Seq cells (left side, Fig. 2b) and Scanorama an unmixed distribution of cells (right side, Fig. 2b). For cases where batch correction largely fails (Supplementary Fig. 7b), though the ‘integrated’ ambient spaces by either method are similar to the pre-integrated ambient space, reduction to 2D can enhance mixing for the ‘integrated’ spaces, but only increase unmixed cells in the pre-integrated space. We found similar problems when the *L*_1_ norm was used and with t-SNE as used in [10] (Supplementary Fig. 6b,c, 7a). Notably, the initial PCA reduction can drive the reversal or distortion of mixing trends, though removal of PCA-coupling does not alleviate this issue (Supplementary Fig. 6c, 7a). Thus it is unclear what patterns of mixing are a result of the efficacy of the integration method, or arbitrary variation introduced by the dimensionality reduction procedure.

A consequence of these findings is that reference mapping procedures, which aim to demonstrate shared structures between batches or datasets, can also result in appearance of false structures (Supplementary Fig. 8). As an example, UMAP has been proposed as a method for transforming or mapping new data given coordinates fit on another dataset [11]. Yet, transforming high dimensional uniformly distributed points with UMAP coordinates from a single-cell dataset imposes a false structure akin to the structure of the original data (Supplementary Fig. 8) (see Supplementary Methods).

### Cluster Validation & Relationships

Beyond the use of dimensionality reduction to “validate” dataset merging, it is common to use two or three dimensional visuals to assess appearances of clusters. This can be to justify or directly generate cluster or cell type assignments [1–3, 14, 15], and to infer properties of clusters (their heterogeneity, separation, or similarity) [6, 13]. Such uses rely on retention of global relationships (Table 1), where local neighbors are less important compared to maintaining group assignment or patterns of separation between groups (Table 1). Distance preservation may also be necessary if conclusions are to be drawn on the extent of separation or locations of clusters (Table 1). However, across datasets of various sizes [37, 48] the prediction of a cell’s type (assignment) based on its neighbors is consistently worse in the 2D embedding space than in higher dimensional representations (even when labels are given as with supervised UMAP, UMAP Sup.) (Fig. 3a) (see Supplementary Methods). This calls into question the added benefit of using such embeddings as validations or representations of cluster assignment.

**Figure 3:**
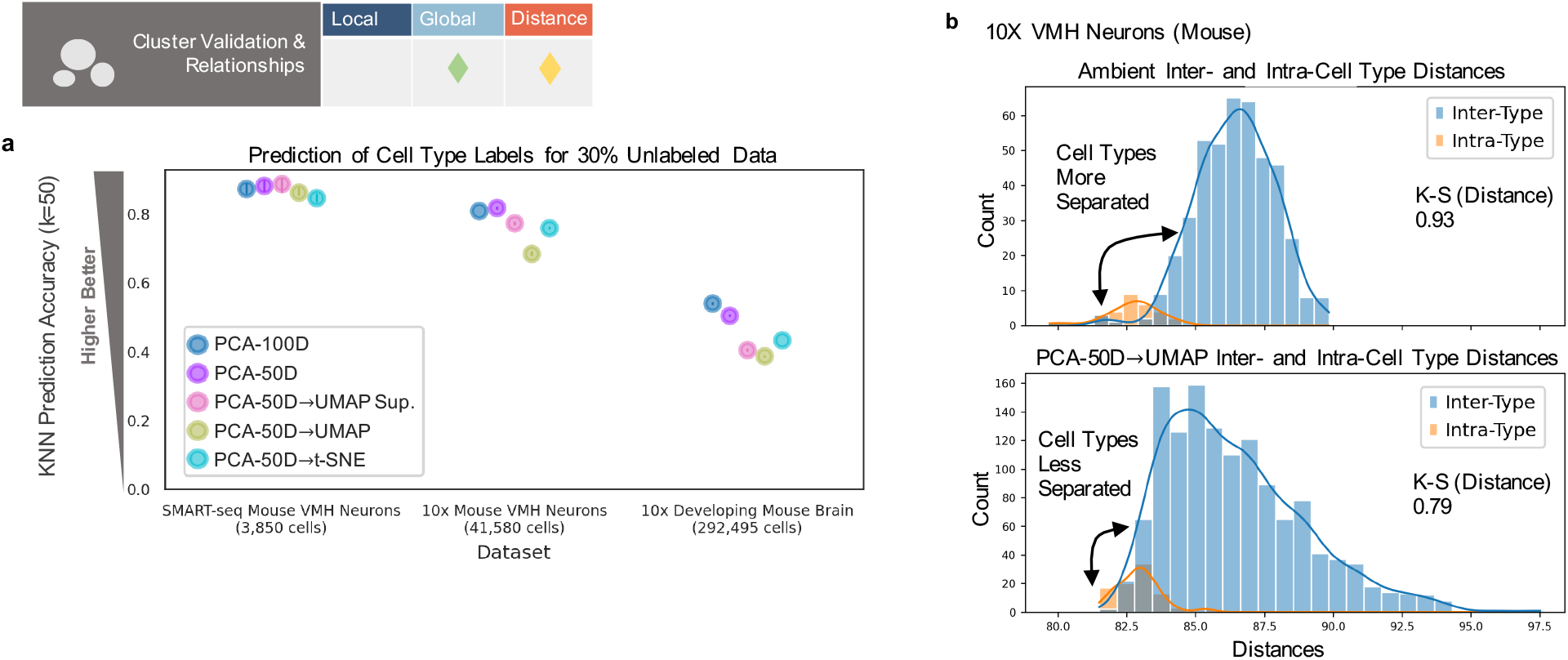
Distortion in Cluster Validation & Relationships. **a)** Prediction of cell type label for 30% of the dataset(s), based on the labels of the 50 nearest neighbors. **b)** Distributions of cell type inter- and intra-type distances for the ambient or reduced space (bottom). K-S distance shown as measure of separation, where higher values denote greater separation (see Supplementary Methods).

Additionally, by comparing the distribution of pairwise distances between cells of different cell types (‘inter-type’) to the distribution of distances between cells within the same types (‘intratype’), we can measure how separated those distributions are i.e. how separated or distinct cell types are from each other (Fig. 3b) (see Supplementary Methods). Though it may be desirable for the low dimensional visualizations to increase separability or clarify cell types as compared to the ambient space, such reduction can have the opposite effect (Fig. 3b), reducing the gap between inter-, intra-type distributions for some datasets and increasing the gap for others, whether using the *L*_2_ or *L*_1_ norm (Supplementary Fig. 9,10).

We found that cluster structures were additionally highly sensitive to the number of neighbors (perplexity for t-SNE) used in constructing non-linear embeddings, a commonly tuned parameter which can range from 1-10% or less of the data [1, 6], in line with other results on the effects of tuning [6, 28]. For the in-utero E10.5 dataset, common choices for this parameter result in different placements and overlaps of cell types, pushing progenitor populations away from their downstream cell states/types or incorrectly merging distinct, early stage populations (Supplementary Fig. 11). Such inconsistencies have led to publication of incorrectly surmised differentiation trajectories from apparent relationships between cell types [49]. Even in a non-biological, machine learning, benchmark dataset [50], we found a muddling of cluster structures, with points belonging to different digits mixed within clusters (possibly hidden by order of points plotted) though high accuracy classification is possible in higher dimensions [51] (Supplementary Fig. 12) (see Supplementary Methods). This reveals a reliance on noise/distortion cancellation to validate cluster assignment and knowledge of ground truth labels to determine the utility of the visual or when tuning is sufficient.

### Density-based Visuals & Marker Analysis

Density assessments of points in 2D embeddings are frequently used to quantitatively assess cell-cell relationships by directly relying on distances between the cells in two dimensions (Table 1). Common applications compare densities of cells in different conditions or batches, within a shared embedding space, to make statements on changes in population density or expression between groups [3, 16, 18, 20]. However, as demonstrated above, parameter tuning easily disrupts the placement of cells and clusters in such visuals, inherently affecting the generation of contours. Furthermore, using different numbers of neighbors for embedding generation can result in dramatic appearances of non-overlapping cell populations (1,4 in Fig. 4a,b) which can disappear when more or less neighbors are used (see Supplementary Methods). Likewise, densities of cell populations can appear the same or different between conditions depending on the number of neighbors used in construction (2,3,5,6 in Fig. 4a,b) (Supplementary Fig. 13,14), confounding the use of these visuals to make comparative statements.

**Figure 4:**
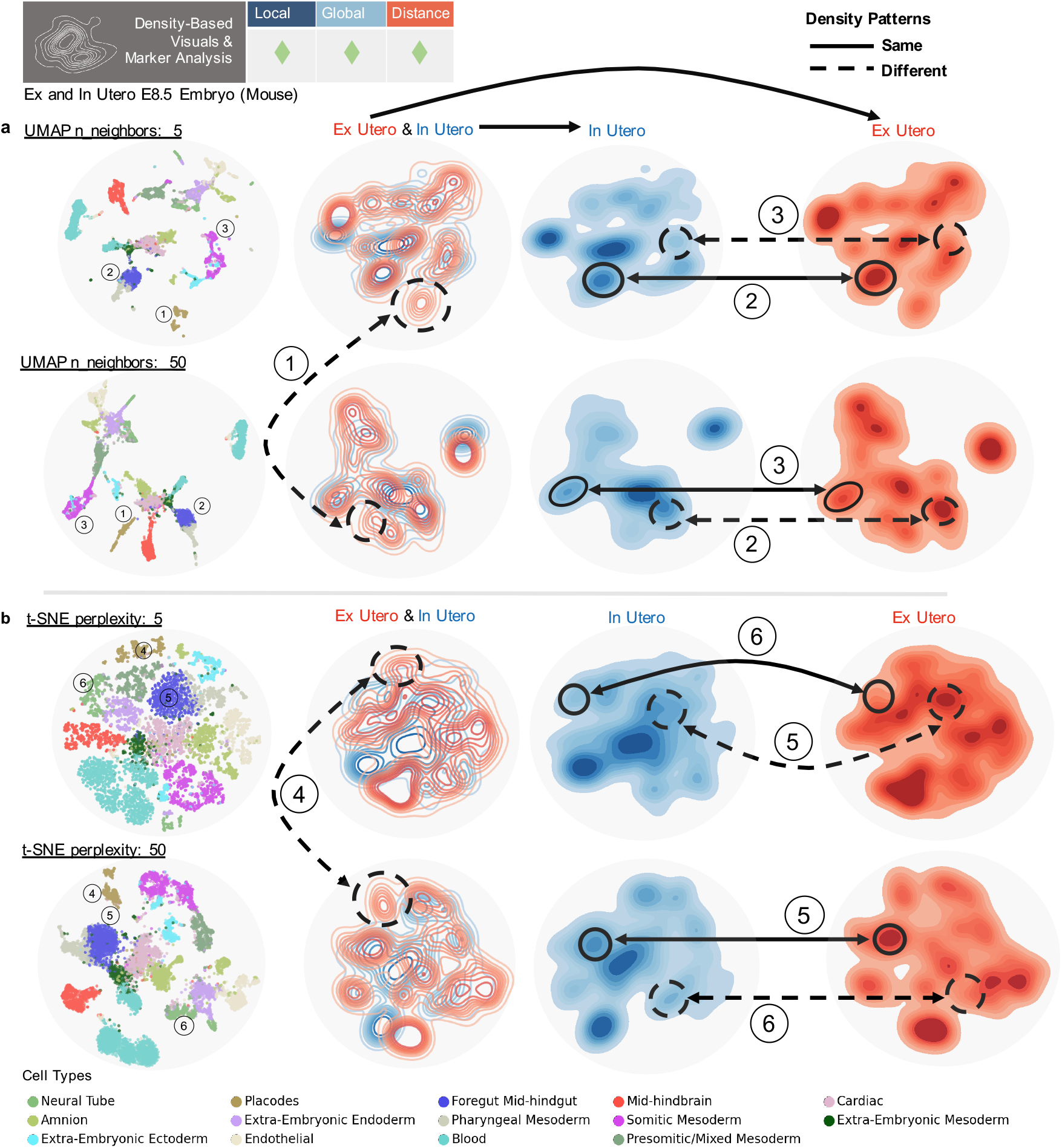
Distortion in Density-based Visuals & Analysis. **a)** Top row (left to right) displays UMAP embedding with n_neighbors = 5, embedding contour plot colored by condition, embedding of just in-utero cells, embedding of just ex-utero cells. Bottom row shows same plots for UMAP embedding with n_neighbors = 50. **b)** Top row shows same plots for t-SNE embedding with perplexity of 5. Bottom row shows same plots for t-SNE embedding with perplexity of 50. Numbers denote comparisons between plots, dashed lines denote a difference, and solid lines denote the same appearance.

### Trajectory Inference & Continuous Relationships

Trajectory inference and pseudotime tasks, such as in RNA velocity [23] or Monocle [22, 24] workflows, focus on local, continuous relationships for inference and calculating pseudotime coordinates. Such tasks may also use distances between embedded points to construct the directions and magnitudes of arrows denoting inferred, developmental trajectories [23, 25] (Table 1). However, as shown with the standard velocyto workflow [23], using the neighbors of cells after reduction to two-dimensions to construct velocity arrows can result in erroneous trajectories, due to the arbitrary placement of cells under different parameter choices. Here we again vary the number of neighbors used to construct the embedding (see Supplementary Methods). Distortions can include loss of continuous relationships, trajectories in incorrect directions, or the addition of new pathways for development (Fig. 5) (Supplementary Fig. 15). Distortions additionally occur due to upstream averaging over nearest neighbors in the inference procedure, and from the choice of embedding procedure (Fig. 5)[52, 53]. Thus the resulting visual compounds distortions from embedding with these prior distortive effects.

**Figure 5:**
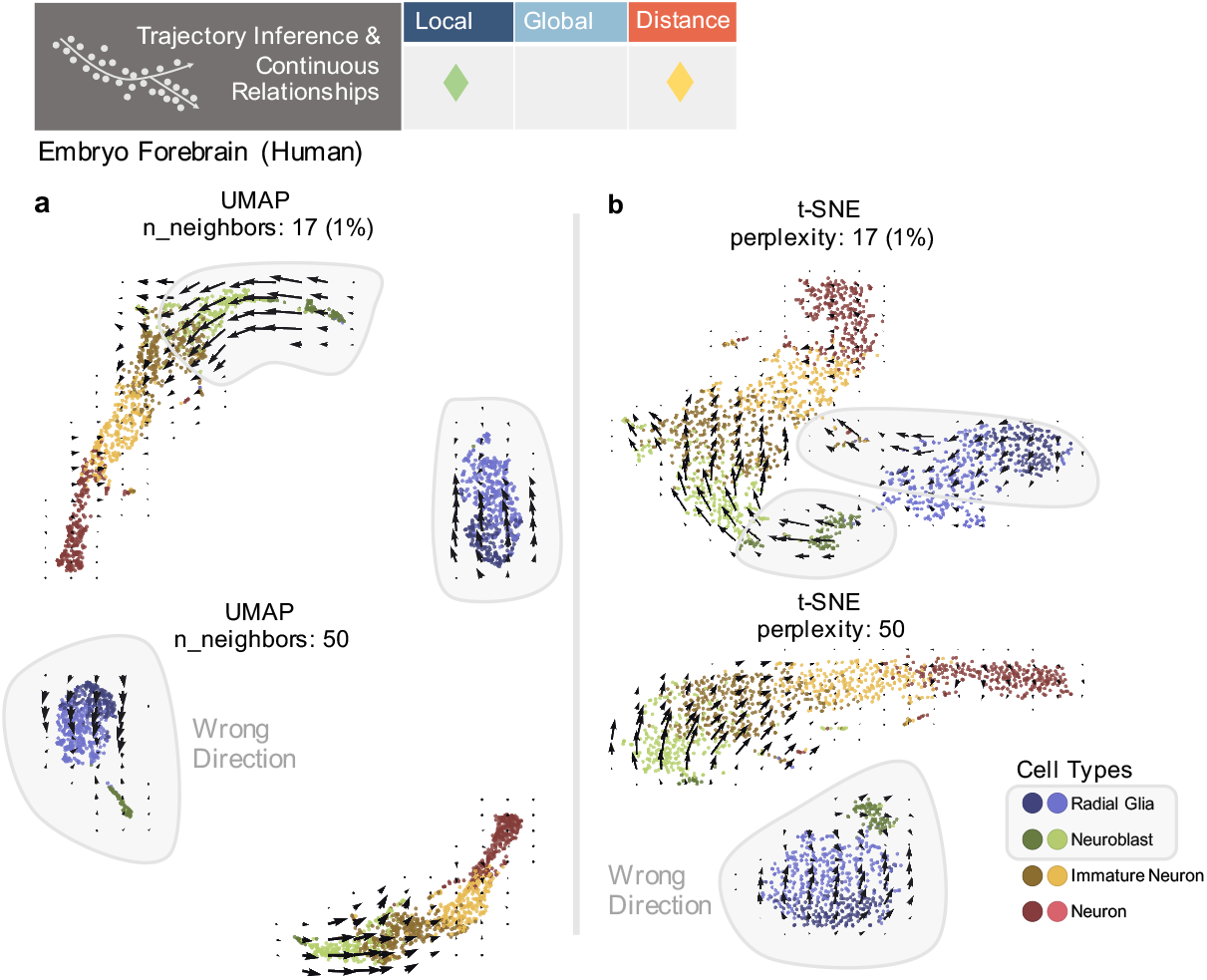
Distortion in Trajectory Inference and Continuous Relationships. **a)** Velocyto RNA velocity embeddings for UMAPs made with 17 or 50 neighbors. Cell types of interest highlighted in grey. **b)** Velocyto RNA velocity embeddings for t-SNEs made with perplexity of 17 or 50.

To investigate distortions of an underlying, continuous manifold by 2D reduction, we used the Swiss-roll as a non-biological benchmark dataset, for which we know the structure in three-dimensions, and moreover is a two-dimensional manifold. We demonstrate how the 3D swiss-roll (constructed by rolling up the two-dimensional plane) loses its coherence when embedded in 2D with UMAP (Supplementary Fig. 16). No embedding recapitulates the original plane [54] and depending on the number of neighbors used, distinct clusters or islands may appear, with a scrambling of local neighbors (made worse by increasing the tightness of the embedded roll) (Supplementary Fig. 16). Thus knowledge of the true manifold is required to understand the disruption of continuity in these embeddings.

Additionally, both cluster-level, global relationships, and locally continuous properties of such visuals are used as independent ‘metrics’ to validate cell type assignment and robustness of clustering results [1, 2, 6, 55]. However, in common single-cell analysis packages (e.g. Scanpy [56] and Seurat [7]), the same k-nearest neighbor (knn) graph constructed from the higher dimensional PCA space is passed to both the clustering algorithm and the embedding algorithm. As shown in Supplementary Fig. 17, the embedding is then not an independent assessment of clustering results and is likely to form clusters that match the knn graph even if that graph does not represent the ‘original’ underlying manifold. Together, the use of such embeddings to imply or infer continuous relationships then becomes an arbitrary endeavour, with a user unable to trust seemingly dramatic connections or isolated populations, and likely to choose what seems most appealing.

### Arbitrary Shapes Preserve Structure

To illustrate the indeterminate nature of two-dimensional UMAP and t-SNE embeddings, we developed an autoencoder framework to fit cells from any dataset to an arbitrary 2D shape, while preserving ambient cell-to-cell distances to an extent not much different than UMAP or t-SNE (see Supplementary Methods) [51, 57, 58]. We found that it is possible to embed data in the shape of a ‘von Neumann elephant’ [59, 60] or a flower. Though it is unlikely scientists would present data in such forms, as shown below, they are equally legitimate in terms of fidelity to the data in ambient dimension, compared to UMAP or t-SNE embeddings. We call this method to produce customized embeddings “Picasso”, in homage to the eponymous artist’s skill in imitating other artistic works.

We compared correlations of inter- and intra-type distances between Picasso embeddings with those of t-SNE, UMAP and PCA, for the ex-utero (E8.5), MERFISH MOp, and SMART-Seq VMH neuron datasets [37]. These distances represent trends often inferred from such visuals, where inter-type distances represent inter-cell-type relationships, and intra-type distances represent the variance or spread within the cell types (see Supplementary Methods). Each Picasso embedding demonstrated comparable performance to t-SNE and UMAP (Fig. 6), even dens-SNE/densMAP [61] (Supplementary Fig. 18), projections, with cells of the same types distinctly grouped together in the arbitrary shapes. Picasso embeddings also improved upon t-SNE/UMAP intra-type correlations for all datasets (Fig. 6). Results were recapitulated for inter- and intra-distances calculated with the L_1_ norm, and for inter- and intra-sex distances for the VMH neuron dataset (Supplementary Fig. 18,19).

**Figure 6:**
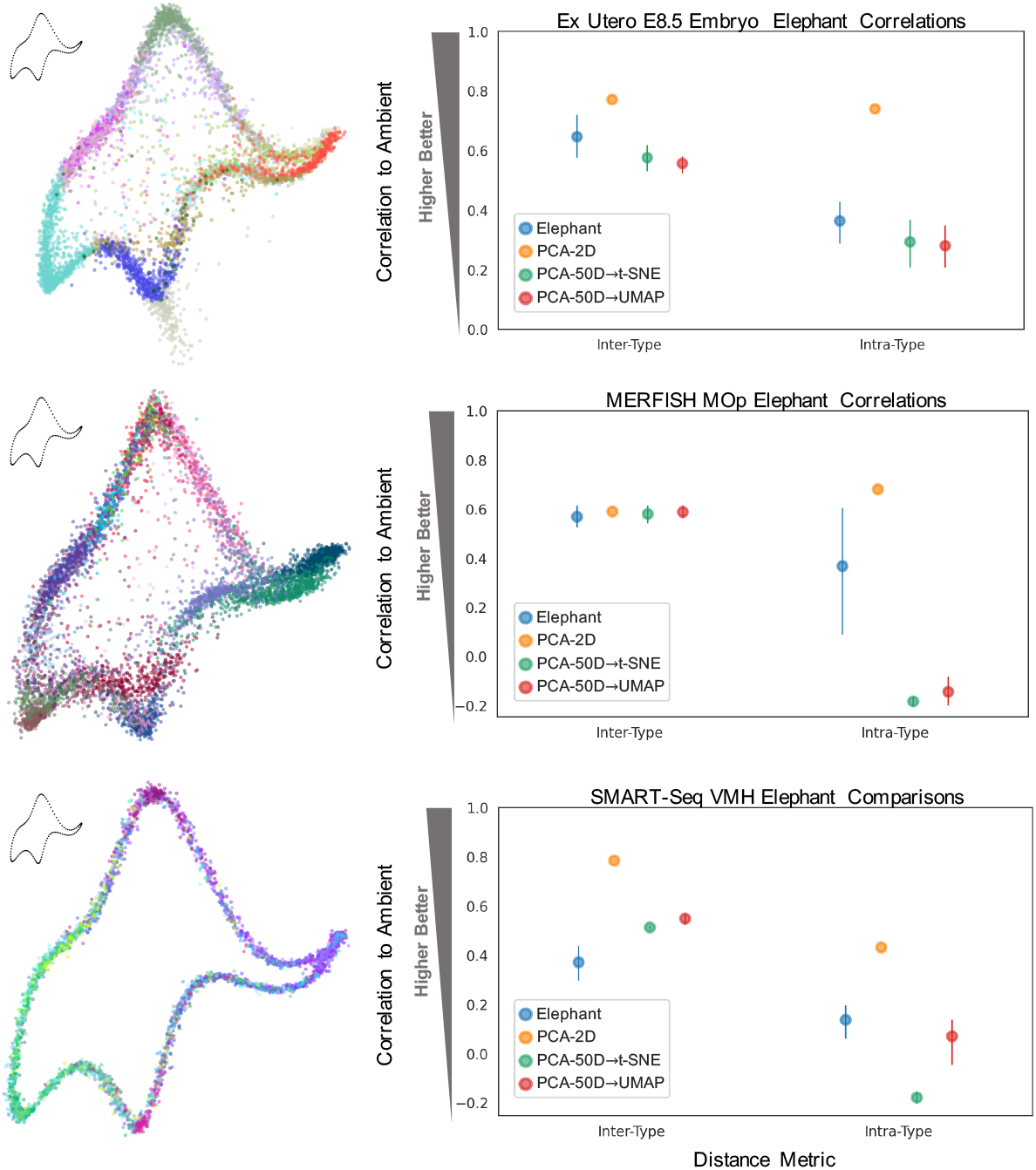
Arbitrary Shapes Preserve Structure. Elephant shape [59, 60] shown on the left, with corresponding correlations of data embeddings to ambient space shown in right-hand plots, for inter- and intra-type distance metrics. Metrics calculated over n=5 embeddings. Colors denote cell types, delineated in Supplementary Fig. 18.

Thus, Picasso can quantitatively represent these visually inferred characteristics similarly to, or better than, the respective t-SNE/UMAP embeddings, while producing arbitrary shapes.

## Final Thoughts and Discussion

### Limitations for Exploratory Data Analysis (EDA)

Although popular two-dimensional embeddings can reflect the broader strokes of the data such as cell type inter-distances, or highlight correlations between features [62], our findings highlight fundamental obstacles in reduction of high-dimensional data to 2D, the generation of multiple, possibly contradictory interpretations of the same data across applications, and the limited utility of these embeddings as EDA tools. Though at the heart of EDA, as defined by statistician John W. Tukey [63–65], is the exploration of data through visualizations prior to confirmatory analysis, such visuals are intended to encompass robust or ‘resistant’ analyses which extract (expected or unexpected) features of the data [63]. Thus the use of these 2D embeddings to reveal expected or unexpected properties is fraught by the fact that it is unclear which properties will be preserved or displayed i.e. the purpose of the visual itself, where seemingly strong characteristics can be arbitrary distortions, from integration/mixing patterns (Fig. 2) to the existence of or connections between clusters (Fig. 3,5, Supplementary Fig. 11). Attempts to show error or significance of cell placement on these visuals do not tackle the inherent limitations of low dimension embedding, how to determine which features are displayed, and what is distortion to ignore [27, 66]. Prior analysis is required to determine ‘sufficient’ parameter tuning and to define the purpose of the visual, undermining the use of such procedures as EDA tools. Together, this results in a user conducting two confounded exploratory analyses, that of the method properties and that of the data properties.

Another of the ‘guiding principles’ of EDA is to conduct “analyses...before summaries” [63], using analyses to present particular features of the data, then to be collated as a summary. However, the use of such all-in-one visuals begins from a place of summary rather than analysis, showing ‘all points and all relationships’ at once and attempting to approximate many properties. In general, the open-ended nature of these visuals and ability of aesthetic parameter tuning to manipulate and create biological patterns demonstrate the ease with which such tools become confirmatory bias aids, and that such 2D spaces should be treated more as cartoon diagrams to be displayed postanalysis. However, in these cases conceptual graphics can be used instead which do not attempt to represent ‘all points and all relationships’ (to avoid over-interpretation), and higher-level diagrams which do not operate at the cell- or point-wise level [67, 68].

### Incoherences in the Dimensionality Reduction Process

The generation of the 2D embedding is additionally a multi-step process, demonstrated here as a coupling of higher dimensional (linear) reduction by PCA to a non-linear reduction to 2D by t-SNE/UMAP. Each step incurs some distortion of the data, where preservation of certain properties by one reduction can be lost by the next, as well as exaggeration of distorted patterns over the steps. However, this procedure is taken as a baseline [6, 28], and there is little discussion of the logic behind this coupling. We do present analyses alternatively measuring distortions with the L1 metric, given its more desirable properties in higher dimensions than Euclidean (L2) distance (see above), but other choices of distance metrics are possible and, whether in ambient or reduced space, can provide different implications and interpretations of the dataset’s properties [33]. In light of this, one might surmise that the non-linear methods instead learn other manifold-specific ‘metrics’ from cell neighborhoods by identifying ‘biological geometries’ (though this is not justified by the original authors [4, 5]). However, methods such as UMAP and t-SNE at their core rely on measuring distances locally, in concordance with common Euclidean analysis methods. For example, this is the case for neighborhood graph construction as used for clustering [69], pseudotime and trajectory inference [21, 70], as well as non-linear embedding (e.g. UMAP/t-SNE) [4, 5]. Notably reduction of the data with PCA prior to non-linear dimensionality reduction is incoherent with the premise that the data lies on a ‘biological manifold’, as PCA implicitly assumes Gaussian noise for data that lies in a Euclidean space. PCA additionally, inherently, reduces variance of projected data, while methods such as UMAP can create variance in embedded data [71].

Utilizing these 2D visuals to infer structure of the underlying manifold then requires knowledge of that manifold itself to interpret these outputs and distortions, a task confounded by noise present in biological data and the fact that common methods poorly recapitulate simple non-Euclidean manifolds (Supplementary Fig. 16) [54]. And while PCA does impose assumptions of Euclidean geometry and Gaussian noise model, the assumptions of heuristic, non-linear methods are more opaque and their results not easily falsifiable.

### Alternative Methods and Analysis Approaches for Applications

We therefore discourage reliance on and blind application of such heuristic procedures, particularly across the range of applications in Table 1. Instead greater focus should be given to utilizing and developing an array of investigative and self-consistent analysis tools, which provide clearer interpretation of their goals and the biological features being assessed, present targeted lowdimensional embeddings and visuals displaying these features, and can easily be combined with statistical procedures to generate and falsify hypotheses.

With respect to the task of representing all neighbors in an embedded space, we demonstrate a use of (semi) supervision to instead balance or preserve *specific* classes of neighbors, particularly given that multiple modalities can co-exist within single-cell data matrices, with each modality represented to a different extent when dimensionality reduction is applied. We utilize higher dimensions for constructing such spaces, to be used for downstream tasks, yet such work could also be extended to create high-level, targeted visuals or cartoons. Adapting the neighborhood component analysis algorithm [72] which performs dimensionality reduction optimizing placement of neighbors of the same label near each other, we develop a multi-class, multi-label (MCML) methodology to balance multiple, desired discrete and/or continuous labels ascribed to cells of the same dataset (e.g. cell type, behavioral condition, pseudotime) (Supplementary Fig. 20). By controlling neighborhood structure, we demonstrate improved prediction capabilities and multi-label transfer/prediction for unlabeled cells in these latent spaces (Supplementary Fig. 20c-e,21) [73]. From the observation that intra-metrics, such as intra-type distances, are less well-preserved in high or low dimensional embeddings, we additionally adapted MCML to bias latent space structure (bMCML) to quantitatively preserve these desired intra-metrics (Supplementary Fig. 22). Such task-oriented latent space construction and label-aware objectives could also be extended to preserve or filter label-based properties which contribute to explaining variance in the data or to the accuracy of a specific task (e.g. spatial location prediction, likely dependent on multiple covariates [74]).

For the applications in Table 1, there are existing methods and metrics, as well as opportunities for method development, which would provide more targeted alternatives in keeping with principles of EDA. For example, the assessment of multi-modal data integration and mixing can be directly calculated, as shown by the metrics in this study, as well as by other metrics on mixing proportions and separation [9] or on the retention of true batch differences (biological variation) [75]. Such analyses can additionally be conducted in the ambient space, which minimizes the distortion/transformation of gene-related properties, useful for downstream experimentation.

For applications regarding clustering, clusters can be generated from higher dimensional embeddings if not from the ambient space itself [34], and given that marker gene expression is the main method of validation of clusters, existing tools such as heatmaps directly display cluster results with the features (genes) which determined these groupings. Dimensionality reduction on the gene space can additionally be used to filter for genes or sets of genes best suited to separating the clusters [76, 77]. By targeting the objective of an embedding in such a manner, one can more directly determine the necessary dimensionality for a given question.

To assess heterogeneity within clusters or relationships between clusters, similarity metrics or distances can be calculated between the cells [33] and displayed with qualitative or quantitative visuals which preserve these metrics, including hierarchical relationship diagrams such as dendrograms and trees [78, 79], or graph-based network diagrams [80, 81]. Higher-level diagrams that do not seek to display all point-wise information can also be used to represent the results of other inter-cluster analyses [68, 82].

Such cluster-level visuals and metrics, as well as metrics on integration and higher dimensional distribution comparisons as presented here, can be used in lieu of analyses based on contour plots generated from 2D coordinates. Regarding trajectories and continuous relationships, higher dimensions should be used to perform inference of differentiation trajectories [21, 68], and incorporation of probabilistic and biophysically-informed inference methods [67, 83, 84], offer falsifiable and interpretable approaches with targeted visualization alternatives.

Though it may seem appealing to produce visuals of ‘all data and all relationships’, common embedding practice distorts data in obscure ways, attempts to pack the capabilities of many different analyses into one space, and is easily manipulated. Given these limitations, and the distortions induced by earlier processing steps [85], it is preferable to limit dimensionality reductions/transformations particularly when the space of interest can be treated directly, to utilize and develop targeted analyses for common questions that enable focused visuals, and collate these analyses to drive downstream, hypothesis-driven biological discovery.

## Data Availability

Download links for the original data used to generate the figures and results in the paper are listed in Supplementary Table 1. Processed and normalized versions of the count matrices are available on [CaltechData](https://data.caltech.edu/), with links provided in Supplementary Table 2.

## Code Availability

All analysis code used to generate the figures and results in the paper is available at https://github.com/pachterlab/CP_2023 with Picasso and MCML analyses provided in notebooks which can be run on Google Colab. Picasso is also available at https://github.com/pachterlab/picasso. The MCML method as well as tools for quantitative analysis are available via a Python pip installable package from https://github.com/pachterlab/MCML.

## Acknowledgements

Some of the computations presented here were conducted using machines in the Resnick High Performance Center, a facility supported by the Resnick Sustainability Institute at the California Institute of Technology. We thank Joeyta Banerjee for her work developing data processing scripts and Colab notebooks for a previous manuscript version [86]. We also thank Gennady Gorin and Benjamin Rivière for helpful discussions regarding the MCML and Picasso analyses, Sina Booeshaghi for helpful discussions regarding NCA and dimensionality reduction, Ingileif Hallgrímsdottir for valuable feedback on the manuscript, and Páll Melsted for useful insights regarding Theorem 1. The work was supported in part by NIH grant U19MH114830.

