## Supplementary Materials for "The Specious Art of Single-Cell Genomics"

### The Specious Art of Single-Cell Genomics
### Supplementary Materials

Tara Chari[1] and Lior Pachter[1,2,*]

[1]Division of Biology and Biological Engineering,
California Institute of Technology, Pasadena, California

[2]Department of Computing and Mathematical Sciences,
California Institute of Technology, Pasadena, California

December 22, 2022

### Supplementary Methods

#### Datasets and Pre-processing

All datasets used in this study are listed in Supplementary Table 1, and were chosen to cover a range of sequencing platforms, experiment sizes, and experimental designs. For certain datasets, links to saved pre-processed matrices (also used in the github Colab notebooks) are provided in Supplementary Table 2.

| Dataset | Technology | Cells | Label Metadata | Download Link |
|---|---|---|---|---|
| Ex and In Utero Mouse Embryo E10.5 | 10x Genomics v3 | 56,528 | Cell Type, Growth Condition | https://ftp.ncbi.nlm.nih.gov/geo/series/GSE149nnn/GSE149372/suppl/ |
| Ex and In Utero Mouse Embryo E8.5 | 10x Genomics v3 | 6,205 | Cell Type, Growth Condition | https://ftp.ncbi.nlm.nih.gov/geo/series/GSE149nnn/GSE149372/suppl/ |
| SMART-Seq Mouse VMH Neurons | SMART-Seq v4 | 3,850 | Cell Type, Sex | https://data.mendeley.com/datasets/ypx3sw2f7c/3 |
| 10X Mouse VMH Neurons | 10x Genomics v2 | 41,580 | Cell Type, Sex | https://data.mendeley.com/datasets/ypx3sw2f7c/3 |
| 10x Developing Mouse Brain | 10x Genomics v1 | 292,495 | Cell Type | http://mousebrain.org/downloads.html |
| Developing C. elegans Embryo (Neural Lineage) | 10x Genomics v2 | 1,075 | Cell Type, Pseudotime | http://staff.washington.edu/hpliner/data/ |
| Mouse Primary Motor Cortex (MOp) | MERFISH | 6,963 | Cell Type, Spatial Coordinates | https://caltech.app.box.com/folder/134209256308 |
| Human Embryo Forebrain | 10x Genomics v1 | 1,711 | Cell Type | https://github.com/tarachari3/GFCP_2022/blob/main/notebooks/data/hgForebrainGlut.loom |
| CEL-Seq Human Pancreatic Islet Cells | CEL-Seq | 1,276 | Technology | http://cb.csail.mit.edu/cb/scanorama/data.tar.gz |
| SMART-Seq2 Human Pancreatic Islet Cells | SMART-Seq2 | 2,989 | Technology | http://cb.csail.mit.edu/cb/scanorama/data.tar.gz |
| inDrop Human Pancreatic Islet Cells | inDrop | 8,569 | Technology | http://cb.csail.mit.edu/cb/scanorama/data.tar.gz |

**Supplementary Table 1: Dataset Metadata.** Datasets used for all distortion and Picasso analyses.

| Name | DOI Link |
|---|---|
| MERFISH MOp | |
| metadata.csv | https://data.caltech.edu/records/2063 |
| counts.h5ad | https://data.caltech.edu/records/2064 |
| 10x VMH Neurons | |
| metadata.csv | https://data.caltech.edu/records/2065 |
| tenx.mtx | https://data.caltech.edu/records/2072 |
| var.csv | https://data.caltech.edu/records/2066 |
| tenx_raw.mtx | https://data.caltech.edu/records/2073 |
| SMART-Seq VMH Neurons | |
| metadata.csv | https://data.caltech.edu/records/2067 |
| smartseq.mtx | https://data.caltech.edu/records/2071 |
| smartseq_raw.mtx | https://data.caltech.edu/records/2070 |
| gene_names.npy | https://data.caltech.edu/records/2068 |
| smartseq.csv | https://data.caltech.edu/records/2075 |

**Supplementary Table 2: Availability of Processed Data.** Links to DOI registered data for any externally pre-processed data used for analyses.

For the SMART-Seq and 10x mouse VMH datasets, cells were filtered according to the steps outlined in [37]. Unless already provided, the top 2000 highly-variable genes (HVGs) were identified for all datasets using Scanpy's highly_variable_genes [56]. Counts were log-normalized, unless previously transformed, with the log-count matrices representing the 'ambient' data for metric comparisons (see below). Thus unless otherwise indicated, 'ambient' space refers to the log-normalized count matrices filtered for HVGs. All count matrices were mean-centered and scaled before application of Picasso or principal component analysis (PCA). All PCA analysis was performed using sklearn TruncatedSVD to 50 dimensions by default. 15 dimensions was used for the PCA of the integrated mouse embryo E10.5 dataset and 100 dimensions for the pancreatic islet datasets, to facilitate direct comparison to the original studies [8,10].

The t-SNE and UMAP algorithms were applied to the higher dimensional PCA embeddings with default settings. This sequence of first PCA reduction, then reduction to 2D, is denoted as 'coupling'. The effect of a single parameter (n_neighbors) change is shown for UMAP embeddings in Fig. 4,5 and Supplementary Fig. 13-17, but we did not adjust parameters beyond this. As per the discussion in [35], though slight changes in parameters can drastically impact low-dimensional embeddings, the choice of parameters for tuning is often informed by empirical observations/prior knowledge leaving open the question of which metric(s) to use for determining 'optimal' parameters. Notably this tuning is also contradictory to the common use or desire of such techniques to produce 'unsupervised' representations of the data.

#### Local Jaccard Distances

For comparisons of nearest neighbor overlaps in PCA and PCA-coupled or non-coupled t-SNE/UMAP spaces we measured the Jaccard distance defined as $1 - \frac{|A \cap B|}{|A \cup B|}$ where $A, B$ represent the sets of each cell's 30 nearest neighbors in the ambient and latent spaces respectively. All embeddings were generated three times to accommodate the non-deterministic nature of these reduction methods.

#### Global Cell Type Neighbor Rankings

To measure preservation of cell type neighbors, we calculated Kendall's Tau correlation of each cell type's neighbor ranking (ranking of which other cell types are its nearest neighbors) to the ambient space rankings, higher dimensional PCA rankings (for coupled embeddings), and between all pairs of UMAPs for Supplementary Fig. 11. We used the average of all $L_2$ (Euclidean) (as in Fig. 1b) or $L_1$ (as in Supplementary Fig. 2) pairwise distances between cells of each cell type to rank the cell type neighbors for each type. As described in the main text we also used the $L_1$ norm for its desirable properties in higher dimensions/transcriptomic applications, and to reduce sensitivity to outliers [29-32,34]. All embeddings were generated three times (n=3).

#### (Equi)Distance Analysis

To find equidistant cells within cell types, we selected cells from within sizeable cell types to narrow the search space, as the algorithm we used, namely clique detection in undirected graphs, is time consuming due to the underlying problem being NP-complete. The cell types we investigated were 'Esr1_6' in the 10x VMH dataset and 'Chondrocytes and Osetoblasts' in the integrated embryo E10.5 dataset. We calculated all pairwise distances between the cells in the ambient (gene expression) space, using the $L_2$ (Euclidean) distance as this is the default metric for determining neighbors in the t-SNE [4] and UMAP [5] algorithms.

Using these pairwise distances we then defined an adjacency graph where two cells were 'adjacent' when their distance fell within certain distance criteria. Given the distribution of distances produced, we defined adjacency in three different ways. We sub-selected for 'near' pairs of cells that all had distances within a half of a standard deviation around the 0.1 quantile mark i.e. all equivalently close to each other. Likewise, for 'far' cells we selected for cell pairs around the 0.9 quantile mark, and for 'mid-range' we selected cell pairs around the mean distance (using the same filtering radius of within a half of a standard deviation). This filtering for small, medium, and large distances helps to limit the size of the search space when looking for cliques of mutually equidistant cells (below), as well as reveal the diversity of equidistant cell-cell relationships.

We used the sklearn pairwise_distances for the pairwise calculations. From the graph of 'connected' cells we looked for cliques, namely subsets of cells in which all cells are connected (adjacent) to each other. To find cliques we used the find_cliques function from the networkx package to detect cliques in undirected graphs.

We used two metrics to assess distortion of equidistant cells in two dimensions. The first is the variance of the pairwise distances between cells (or centroids) in each group, as compared to the variance of the distances in the ambient space. We also calculated the ratio of the maximum to minimum distance between cells in each group (the 'max/min ratio'), a quantity for which we derived a lower bound (see Theorem 1 in the Supplementary Note) :

$$\frac{D}{d} \geq \sqrt{\frac{n-2}{2}}.$$

All variance and min/max comparisons were done in the ambient space, the PCA-reduced spaces, and the UMAP/t-SNE spaces, generated with and without coupling to the higher dimensional PCA-reduced space. The ambient space for the integrated embryo E10.5 data is the 'Variance-Stabilized and Scaled' data (as opposed to solely 'Log-Normalized' counts), as this was used as input for the original UMAP embedding in [8], and accompanies the analysis in Fig. 1c, 2a.

These distortion metrics were also measured between every cell and its 10 nearest neighbors to demonstrate distortion outside of groups of necessarily equidistant cells. The sklearn NearestNeighbors function was used to find these 10 neighboring cells for max/min ratio calculations in the embedded versus ambient spaces.

#### Mixing Analysis

All calculations to asses mixing were replicated with the $L_2$ and $L_1$ metrics. To assess mixing for the integrated E10.5 data and pancreatic islet cell datasets, the fraction of each cell's 30 nearest neighbors in its same condition/batch was calculated in the ambient and reduced spaces.

We used the Scanorama correct() [10] and MNN mnnCorrect()[47] methods to batch-correct the datasets in Fig. 2 after highly variable gene selection, then performed standard PCA and non-linear 2D reduction.

#### Metrics for Cluster Relationships

For testing prediction/classification capabilities of embeddings, we used the 50 nearest neighbors of the designated 'unlabeled' cells (30% of the dataset) to determine its cell type label. We implemented this with the sklearn KNeighborsClassifier. For UMAP Supervised (UMAP Sup.), see https://umap-learn.readthedocs.io/en/latest/supervised.html, we provided labels for 70% of the cells.

For inter- and intra-type distances we use the inter-type distances from the cell ranking analysis above (i.e. average pairwise distance between cells of the different types) and calculate intra-type distances as the average pairwise distance between cells within each type). To provide a quantitative measure on the separation of these distance distributions we use the two-sample Kolmogorov-Smirnov test statistic with higher values indicating greater separation (less overlap).

To assess malleability of cluster structures the log-normalized in-utero E10.5 data was reduced to 50D with PCA and projected to 2D with UMAP. Only the n_neighbors parameter was changed, as this is the most commonly tuned parameter.

For the MNIST dataset, we embedded the data into two dimensions without PCA-coupling as that is not standard for non-biological data. 'Color Controlled' plots were generated by plotting the majority cell type in each k-means cluster first then plotting the cells that were obscured on top. We also used the k-means function from sklearn to determine clusters (where k is the number of desired digits) from the two-dimensional embeddings, and measured the digit neighbor rankings and intra-digit correlations as described for the inter- and intra-type calculations above.

#### Density-Based Analysis

For the density/contour plot analysis we embedded the log-normalized and integrated ex- and in-utero E8.5 dataset varying the n_neighbors (perplexity) parameter for UMAP or t-SNE. Contours were generated using the kdeplot function from seaborn.

#### Trajectory Analysis

Using the velocyto package, we generated the RNA velocity embedding for the forebrain dataset used in [23], following Fig. 7 in [52]. Only the n_neighbors parameters were varied for the final, 2D embedding step.

#### Picasso Embeddings & Metrics

The autoencoder network used in the Picasso algorithm is outlined below. The input is a centered/scaled count matrix $\mathbf{X} \in \mathbb{R}^{n \times g}$, $n$ cells by $g$ genes. The input is passed through two fully-connected layers of 128 nodes and $d$ nodes respectively with $d = 2$ by default. Batch normalization, the ReLU activation function, and dropout regularization are applied between the layers. The second layer represents the latent representation in $\mathbb{R}^{n \times d}$ denoted as $\mathbf{Z}$. The final linear, decoder layer produces $\hat{\mathbf{X}} \in \mathbb{R}^{n \times g}$. No activation function or bias terms are used between the latent and decoder layer as the decoder output solely represents a linear transform of the latent space.

Mini-batch training was employed, with a default batch size of 128, though larger batch sizes were used for some Picasso embeddings. Adam optimization [58] was used for network training with a default learning rate of $10^{-3}$ and weight-decay term of $10^{-5}$.

We defined two loss functions: $L_{ShapeAware}$ and $L_{Reconstruction}$, which balance the fit of the input points to the desired shape coordinates and reconstruction error in the decoder output as compared to the input. $\mathbf{S} \in \mathbb{R}^{p \times d}$ represents the coordinates comprising the desired shape, where $d = 2$ and $p \geq n$. The latent space $\mathbf{Z}$ is also limited to $d = 2$ dimensions. The pairwise distance matrix $\mathbf{D} \in \mathbb{R}^{n \times p}$ represents Euclidean distances between the cell coordinates in $\mathbf{Z}$ and shape coordinates $\mathbf{S}$ such that

$$d_{ij} = \|z_i - s_j\|_2.$$

Using $\mathbf{D}$, we define a Boolean, $n \times p$ adjacency matrix $\mathbf{A}$, where $\sum A_i = 1$. This matrix uniquely specifies an adjacent coordinate point for every cell, in a bipartite graph mapping the $n$ cells to the $p$ coordinates. $\mathbf{A}$ is determined by the linear_sum_assignment SciPy package, which assigns a shape coordinate to each cell by solving the minimization problem:

$$min \sum_i \sum_j d_{ij} a_{ij}$$

where $a_{ij} = 1$ iff row $i$ is assigned to column $j$. Thus,

$$L_{ShapeAware} = \sum A \odot D.$$

Picasso performs this minimization to attempt to map cells to their closest, unique shape coordinates. The reconstruction loss is the $L_2$ norm of the difference between the reconstructed and input data:

$$L_{Reconstruction} = \|\hat{\mathbf{X}} - \mathbf{X}\|_2.$$

The total loss then incorporates both loss functions, balancing their contributions with $f$, a user-defined fraction weighting the effect of each term on the resulting embedding:

$$L = f * L_{ShapeAware} + (1 - f) * L_{Reconstruction}. \tag{1}$$

The inter-type (or inter-sex) distances were calculated as the distances between the type centroids, or average distances between cells of each sex within each type. Intra-type (or intra-sex) distances were calculated as the average pairwise distance between cells within the types, or within the sexes within each type. Both $L_2$ and $L_1$ distance metrics are provided for these analyses. Pearson correlation was reported between these distances in the embedded spaces and the corresponding ambient space.

#### MCML Embeddings & Metrics

'MCML' (multi-class multi-label) denotes the semi-supervised, label-aware methodology which directly incorporates the label-aware cost into the latent space structure. Similar to Picasso, MCML uses an autoencoder network with a centered/scaled count matrix $\mathbf{X} \in \mathbb{R}^{n \times g}$, $n$ cells by $g$ genes, as input. For MCML embeddings $C$ is the set containing label vectors for each class $k$, $C : \{\mathbf{c}_1, ..., \mathbf{c}_k\}$. Classes can be discrete or continuous, and multi-dimensional in the case of continuous classes (e.g. cell type, sex, condition, location).

The input is passed through two fully-connected layers of 128 nodes and $d$ nodes respectively with $d = 50$ by default. Batch normalization, the ReLU activation function, and dropout regularization are applied between the layers. The second layer represents the latent representation in $\mathbb{R}^{n \times d}$ denoted as $\mathbf{Z}$. The final linear, decoder layer produces $\hat{\mathbf{X}} \in \mathbb{R}^{n \times g}$. No activation function or bias terms are used between the latent and decoder layer as the decoder output solely represents a linear transform of the latent space, for if such interpretability is desired. Mini-batch training was employed with a default batch size of 128. Adam optimization was used for network training with a default learning rate of $10^{-3}$ and weight-decay term of $10^{-5}$.

For MCML we used two loss functions: $L_{LabelAware}$ and $L_{Reconstruction}$, where $L_{Reconstruction}$ is as defined in (1). For $L_{LabelAware}$, we utilized the Neighborhood Component Analysis (NCA) algorithm from [72]. For all cells a pairwise probability matrix $\mathbf{P} \in \mathbb{R}^{n \times n}$ was created where

$$p_{ij} = \frac{exp(-\|z_i - z_j\|^2)}{\sum_l exp(-\|z_i - z_l\|^2)} \ , \ \sum_j p_{ij} = 1.$$

For discrete labeled data (e.g. cell type names) we defined $L_{Discrete}$ for all pairs of cells $i, j$ where

$$L_{Discrete} = \sum_k \frac{\sum_{ij} p_{ij} \mathbb{1}_{ij}}{\sum_{ij} \mathbb{1}_{ij}} \text{ where } \mathbb{1}_{ij}(\mathbf{c}_k) := \begin{cases} 1 & \text{if } \ c_{k,i} = c_{k,j} \ , \\ 0 & \text{otherwise } . \end{cases}$$

Only the probabilities of cell pairs which are of the same label, for each class $k$, were summed and normalized to the total number of these cell pairs (which represents the maximum value of the numerator). For continuous classes of labels, such as spatial coordinates or pseudotime values, we used a separate loss function, $L_{Continuous}$. A probability weight matrix $\mathbf{W} \in \mathbb{R}^{n \times n}$ was generated for every pair of cells such that

$$w_{ij} = \frac{exp(-\|c_{k,i} - c_{k,j}\|^2)}{\sum_l exp(-\|c_{k,i} - c_{k,l}\|^2)} \ , \ \sum_j w_{ij} = 1.$$

In place of the indicator function, the weights biased the masking of the original probability matrix $\mathbf{P}$ towards closer pairs of cells. Probabilities were also normalized to the maximum of the numerator (treating the weights $\mathbf{W}$ as constants):

$$L_{Continuous} = \sum_k \frac{\sum_{ij} w_{ij} p_{ij}}{\sum_i max(\mathbf{w}_i)}.$$

The final loss function was

$$
\begin{aligned}
L_{LabelAware} &= L_{Discrete} + L_{Continuous} \\
L &= -f * L_{LabelAware} + (1-f) * L_{Reconstruction}.
\end{aligned} \quad (2)
$$

$L_{LabelAware}$ was negated for minimization and was additionally weighted by a constant factor of 10 in comparison to $L_{Reconstruction}$. This factor can be additionally optimized via a parameter grid-search.

For comparisons between the MCML and sklearn's NCA, we ran both methods on the 10x VMH neuron and MERFISH MOp datasets, and MCML was run with $f = 1$ (no reconstruction error) and sklearn's NCA with default settings, to produce 50 dimensional latent space representations incorporating cell type labels only. The NCA loss [72], represented by $L_{Discrete}$, was measured for the generated latent spaces ('NCA Likelihood').

To assess label transfer/prediction in MCML spaces compared to standard embedding spaces (PCA and LDVAE [57]) and SCANVI [73], a common annotation/label transfer method), we used MCML to balance both sex (female/male) and behavioral condition (e.g. aggression, mating, not receptive) of the animals used in the 10x VMH neuron dataset. We measured neighborhood structure in these emebddings by the fraction of cells' neighbors in the same label (for the condition labels or the sex labels), as with the mixing metrics above. Higher fractions (close to 1) denote neighbors all within the same label, as desired. We used the KNClassifier described above to predict sex and condition labels for 20% unlabeled cells, with SCANVI only provided the condition label (as it is not clear how designate multiple categories of labels). For prediction using SCANVI, we used their default reference mapping approach outlined here https://docs.scvi-tools.org/en/stable/tutorials/notebooks/scarches_scvi_tools.html. MCML Ref. utilizes a reference mapping approach akin to SCANVI, where labeled data is embedded, and nearest neighbors labels are then designated for the query (unlabeled) data. However, MCML can also embed all the data together (MCML Full), with only partial labels given for the subset of labeled cells (i.e. using semi-supervision).

We utilized the MCML Full approach to embed the *C. elegans* developing neurons dataset and MERFISH MOp data with continuous labels, pseudotime coordinates or spatial coordinates respectively. We used Jaccard distance to measure the retention of these continuous neighbors in the embedded spaces, for the unlabeled cells. We then generated embeddings with cell type only, cell type and spatial labels, or spatial only and used the KNClassifier to determine accuracy of prediction of both features for unlabeled cells.

To extend this to bMCML (biased MCML), we simplified the targeted reconstruction loss to utilize only one term. Here $L$ is defined by the Pearson correlation of the inter- or intra-distances (see below) of a particular class to the ambient data. $\mathbf{X}$, $\mathbf{b}$ represents the vector of the specified inter-/intra-distances in the ambient space and $\hat{\mathbf{b}}$ represents those same distances calculated for the reconstruction $\hat{\mathbf{X}}$.

$$L = -\frac{\sum_i (\hat{b}_i - \bar{\hat{b}})(b_i - \bar{b})}{\sqrt{\sum_i (\hat{b}_i - \bar{\hat{b}})^2 (b_i - \bar{b})^2}}. \tag{3}$$

#### Supplementary Figures

##### Distortion of Necessary Properties

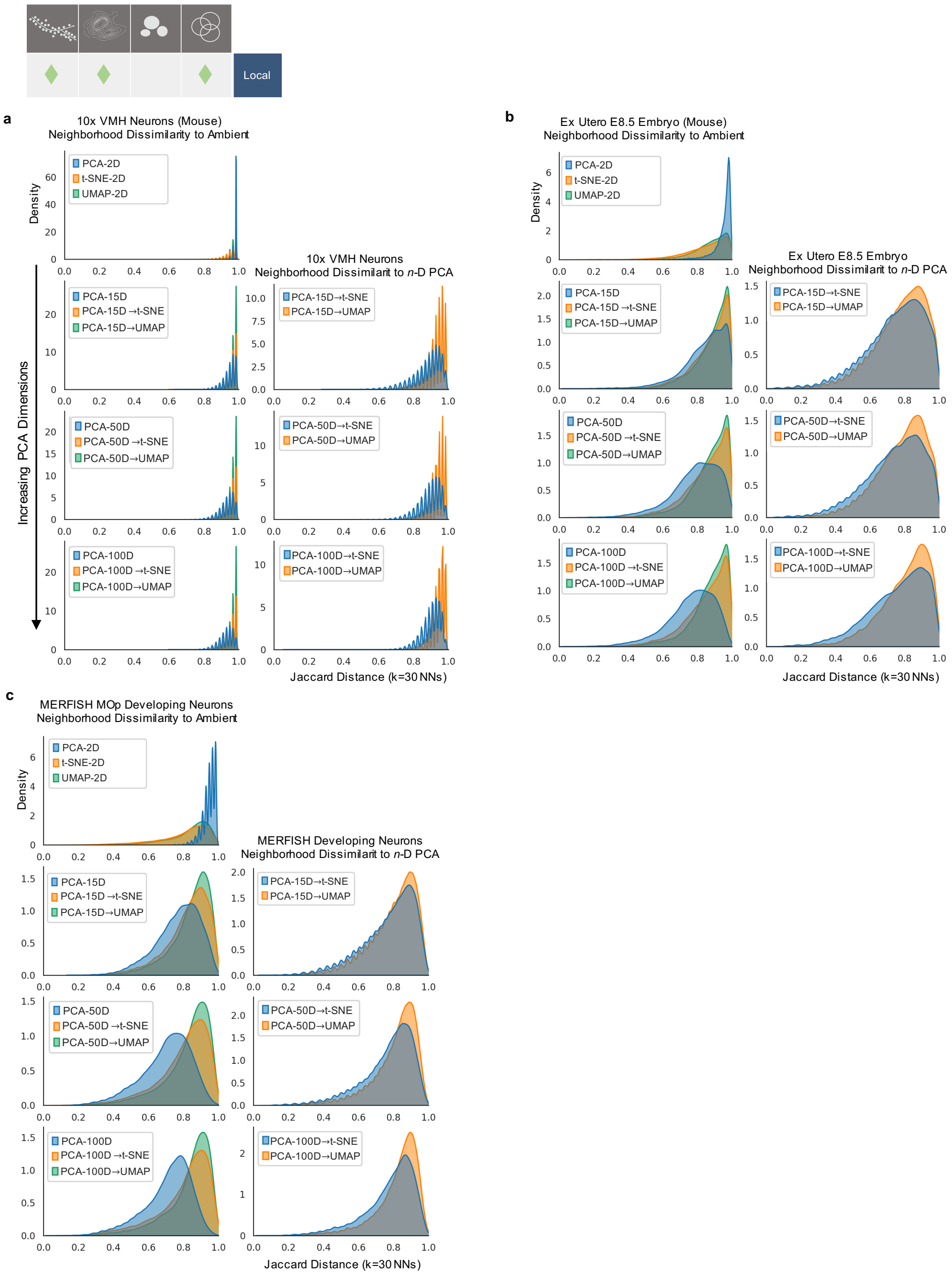

**Supplementary Figure 1: Jaccard Distance Across Datasets a)** Left column, Distributions of Jaccard distances of neighbors compared the ambient data, for the 10X VMH neurons, for uncoupled or coupled 2D embeddings, with increasing PCA dimensions for coupled embeddings down the column. Right column, Distributions of Jaccard distances of neighbors compared the higher dimensional PCA embeddings, for the 10X VMH neurons, for uncoupled or coupled 2D embeddings. **b)** Left and right columns as defined before, with Jaccard distance distributions for the ex-utero E8.5 dataset. **c)** Left and right columns as defined before, with Jaccard distance distributions for the MERFISH MOp dataset.

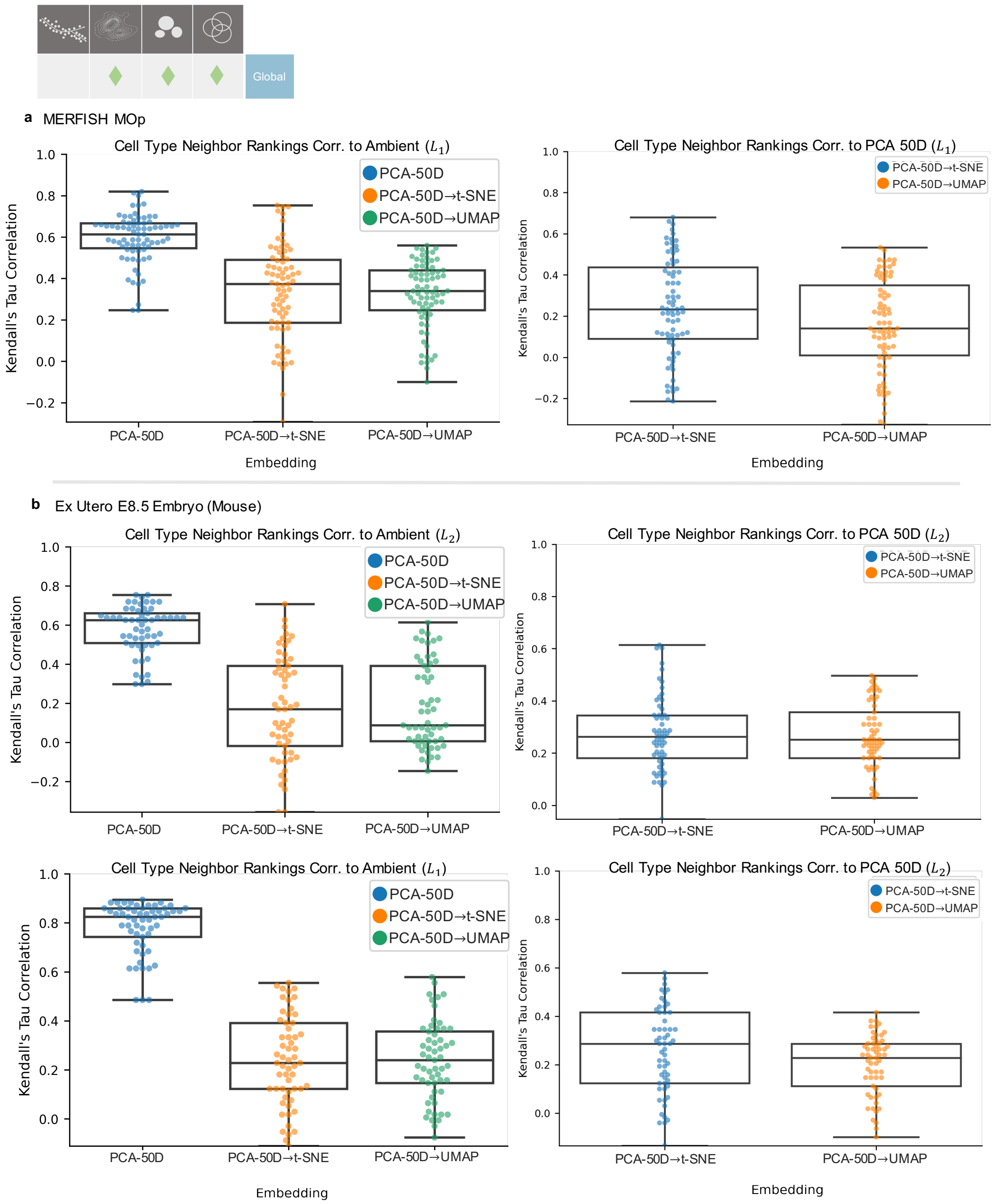

**Supplementary Figure 2: Cell Type Rankings Across Datasets a)** (Left) Kendall's Tau correlation of cell type neighbor rankings (of MERFISH MOp embedded data) to ambient data. (Right) Kendall's Tau correlation of cell type neighbor rankings (of MERFISH Mop embedded data) to the higher dimensional PCA embedding. Rankings calculated with $L_1$ distance metric instead of $L_2$ as shown in Fig. 1b. **b)** (Top Left) Kendall's Tau correlation of cell type neighbor rankings (of ex-utero E8.5 embedded data) to ambient data. (Top Right) Kendall's Tau correlation of cell type neighbor rankings (of ex-utero E8.5 embedded data) to the higher dimensional PCA embedding. Rankings calculated with $L_2$ distance metric. (Bottom Left, Right) Same plots as 'Top' plots with rankings calculated with $L_1$ distance metric. Whiskers denote 1.5 times the IQR. Plots for n=3 different rounds of embeddings.

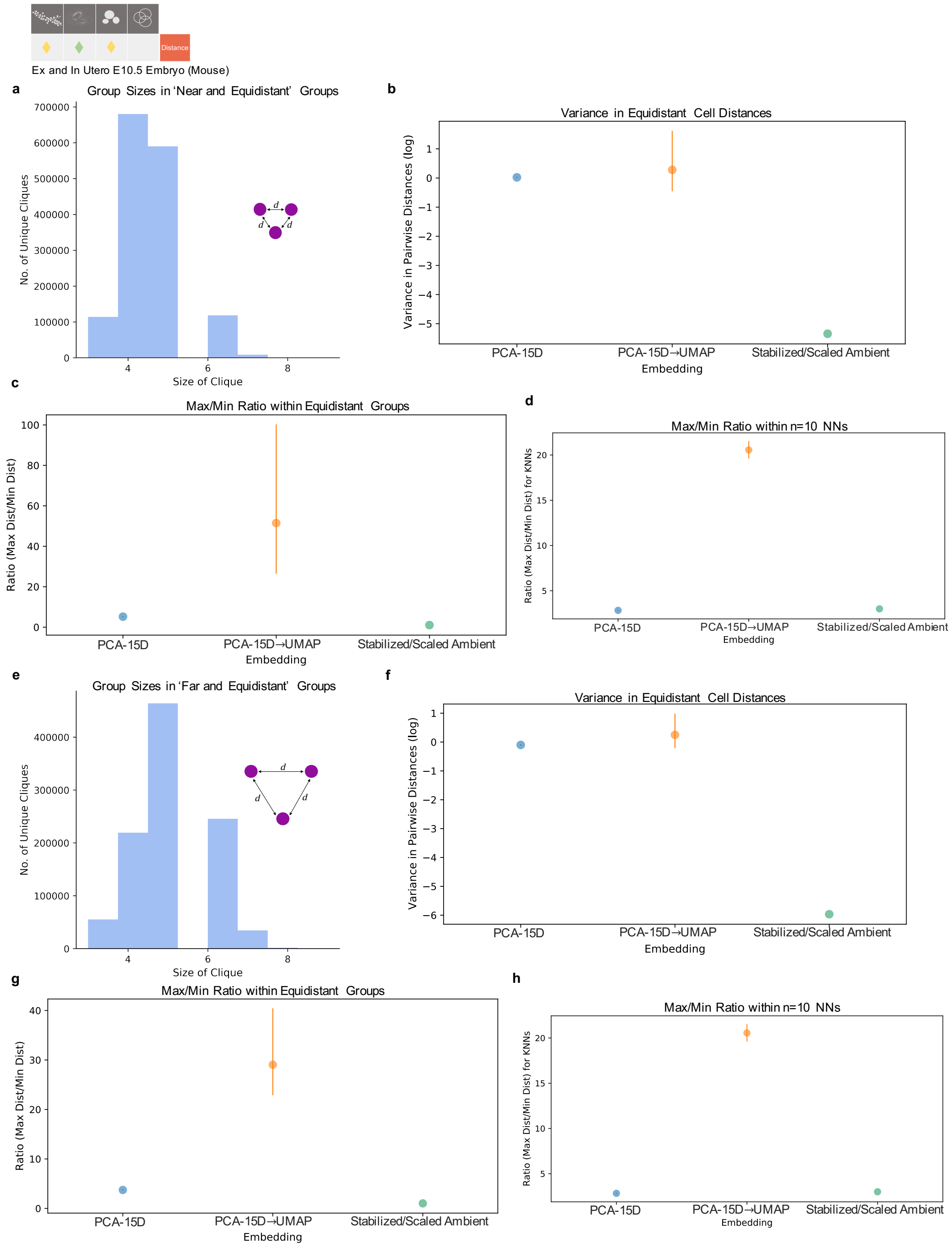

**Supplementary Figure 3: Embeddings of Near and Far Equidistant Points in Integrated Ex- and In-Utero E10.5.** **a)** Histogram of group sizes (number of equidistant cells) in the selection of 'near and equidistant' groups. **b)** Variance of pairwise distances across groups in each latent space. **c)** Ratio of the maximum to minimum pairwise distance (max/min ratio) across groups. **d)** Ratio of the maximum to minimum pairwise distance (max/min ratio) for each cell's neighborhood of 10 nearest neighbors (NNs). **e)** Histogram of group sizes (number of equidistant cells) in the selection of 'far and equidistant' groups. **f)** Variance of pairwise distances across groups in each latent space. **g)** Ratio of the maximum to minimum pairwise distance (max/min ratio) across groups. **h)** Ratio of the maximum to minimum pairwise distance (max/min ratio) for each cell's neighborhood of 10 nearest neighbors (NNs). For all plots bars denote the 95% C.I. and were run over 3 rounds of generated embeddings.

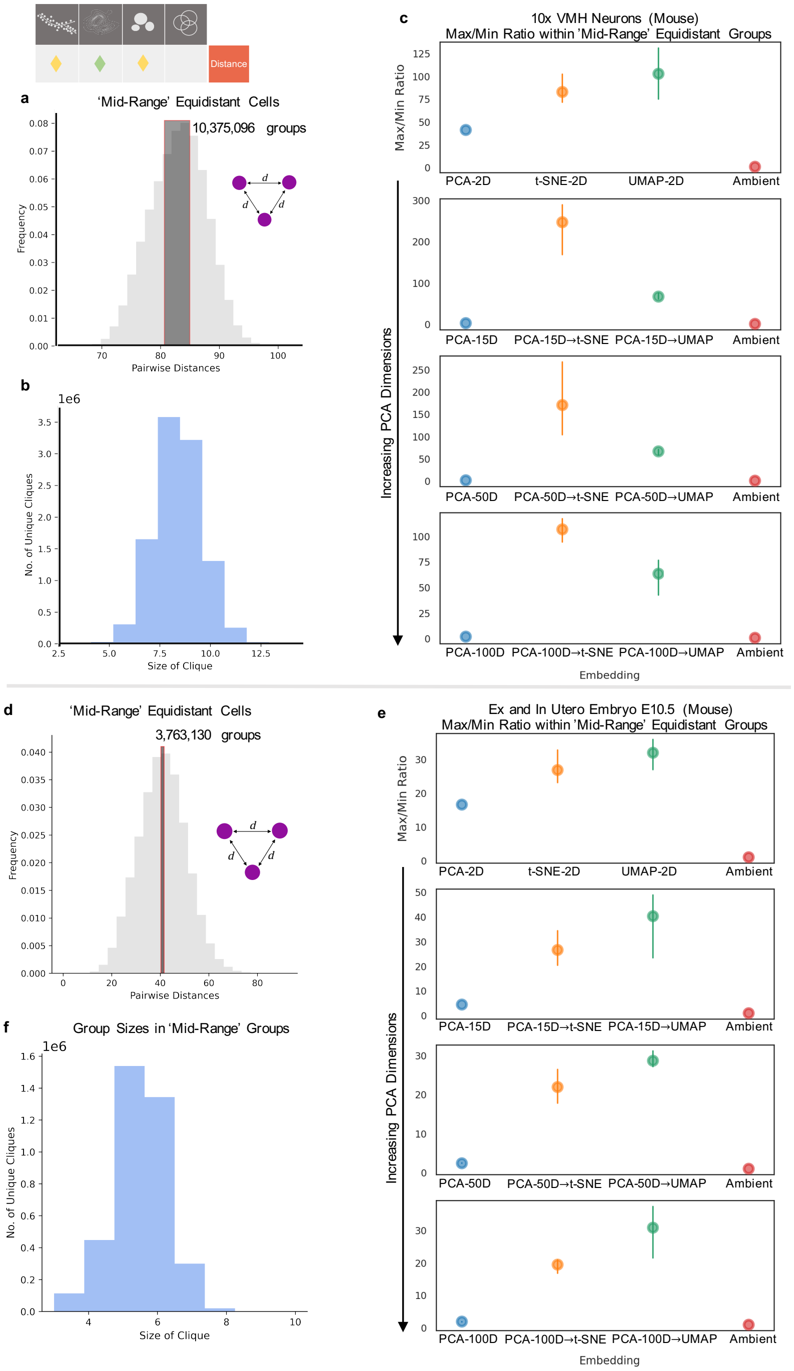

**Supplementary Figure 4: Properties of Equidistant Points in 10x VMH Neurons and Integrated E10.5 Data.** **a)** Selection of 'mid-range' groups, with distances close to the average pairwise distance, in the 10x Mouse VMH Neurons dataset. **b)** Histogram of group sizes (number of cells in a group) in the selection of 'mid-range' groups. **c)** Ratio of the maximum to minimum pairwise distance (max/min ratio) withins groups in latent spaces. Latent spaces with increasing PCA dimensions (if coupled) down the y axis. **d)** Selection of 'mid-range' groups, with distances close to the average pairwise distance, in the Integrated E10.5 dataset. **e)** Histogram of group sizes (number of cells in a group) in the selection of 'mid-range' groups. **f)** Ratio of the maximum to minimum pairwise distance (max/min ratio) within groups in latent spaces.

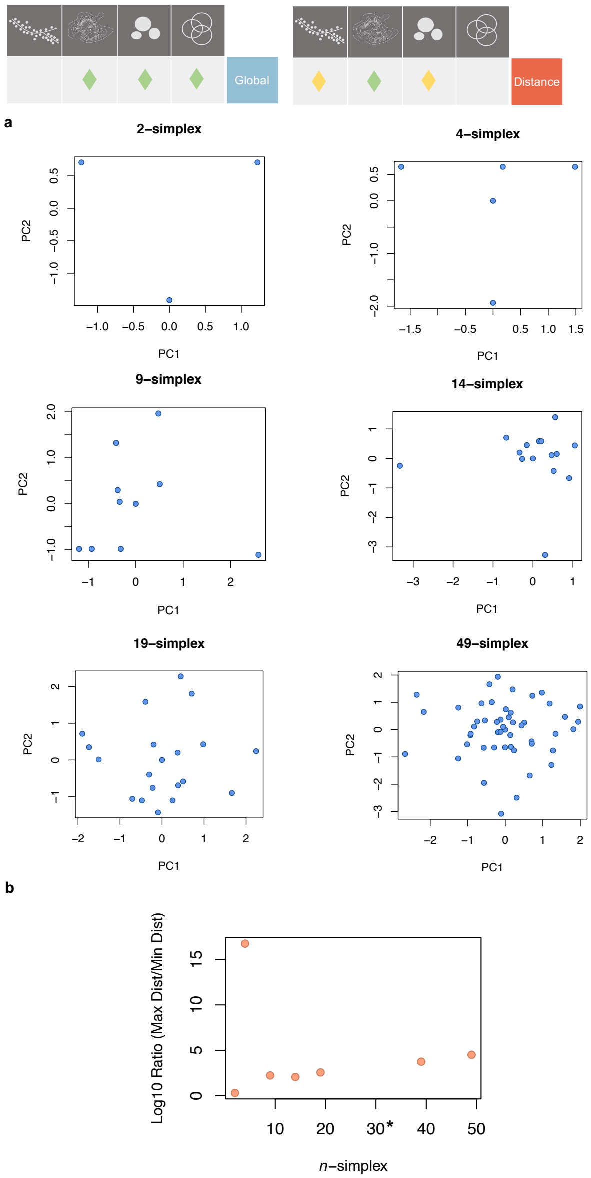

**Supplementary Figure 5: Principal Components of Equidistant Points. a)** First and second principal components shown for varying numbers of equidistant points, i.e. the $\mathbf{I}^n$ identity matrix in $\mathbb{R}^n$, for $n = 3, 5, 10, 15, 20$ and 50. **b)** Max/min ratios for the projections (see Supplementary Methods ; Supplementary Note) of simplices in two-dimensions. * denotes where the minimum distance in the ratio is 0 (points are collapsed onto each other), and the max/min ratio is infinite.

### Distortion in Applications

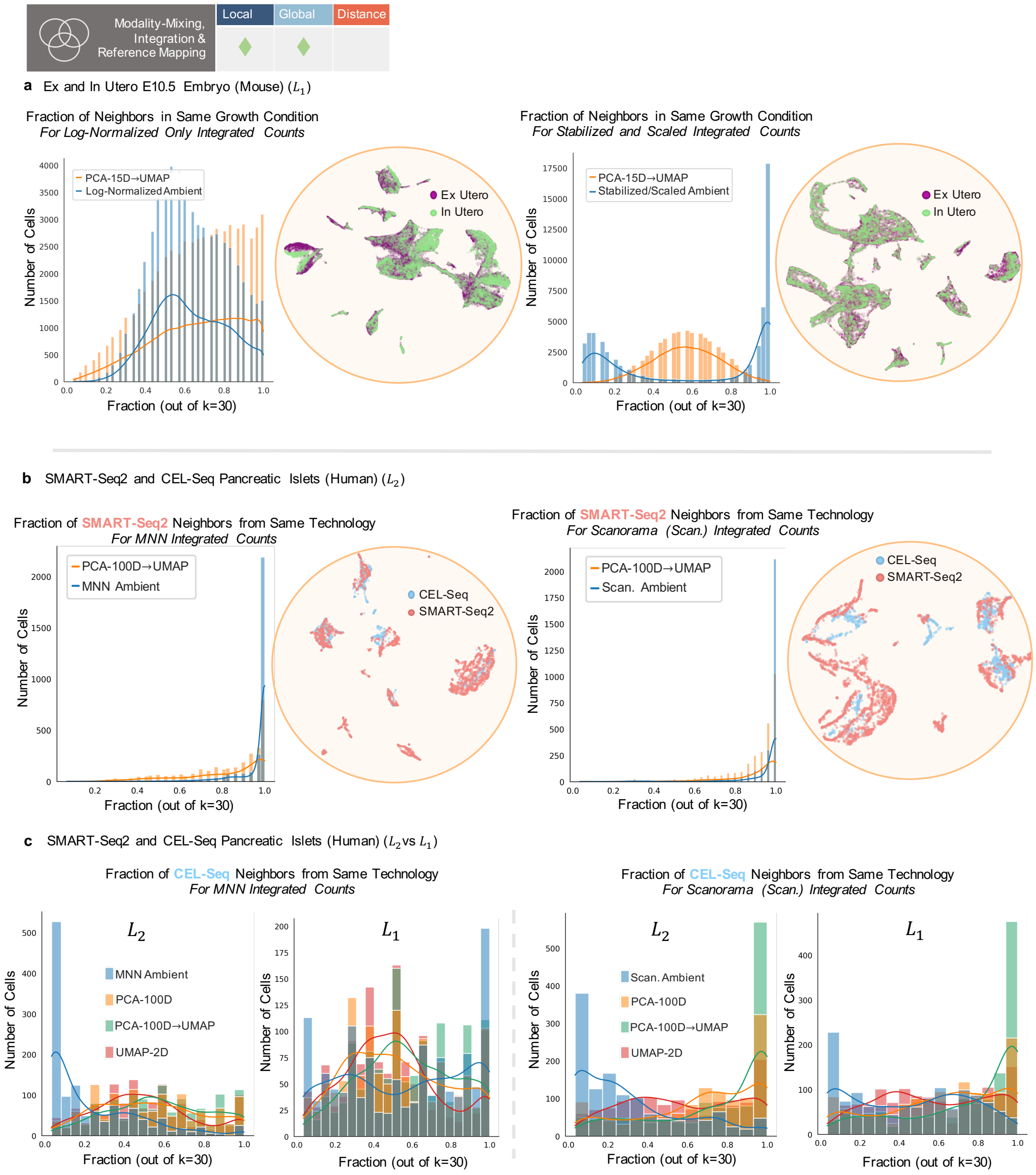

**Supplementary Figure 6: $L_1$ Analysis of Mixing Patterns.** **a)** Left plot shows 'Log-normalized' ambient (blue) and embedding (orange) distributions of mixing (fraction of cell neighbors in the same condition), where 1.0 is no mixing. Corresponding UMAP embedding shown next to it. Right plot shows 'Variance Stabilize and Scaled' ambient (blue) and embedding (orange) distributions of mixing (fraction of cell neighbors in the same condition). Corresponding UMAP embedding shown next to it. Neighbor determination done with $L_1$ distance. **b)** Left plot shows 'MNN Integrated' ambient (blue) and embedding (orange) distributions of mixing (fraction of cell neighbors in the same condition) for SMART-Seq2 cells. Corresponding UMAP embedding shown next to it. Right plot shows 'Scanorama Integrated' ambient (blue) and embedding (orange) distributions of mixing (fraction of cell neighbors in the same condition) for SMART-Seq2 cells. Corresponding UMAP embedding shown next to it. Calculations done with $L_2$ distance. **c** Comparison of neighbor fraction/mixing distributions for CEL-Seq cells calculated with $L_2$ or $L_1$ distance. Distributions shown for all intermediate embeddings and UMAPs without coupling.

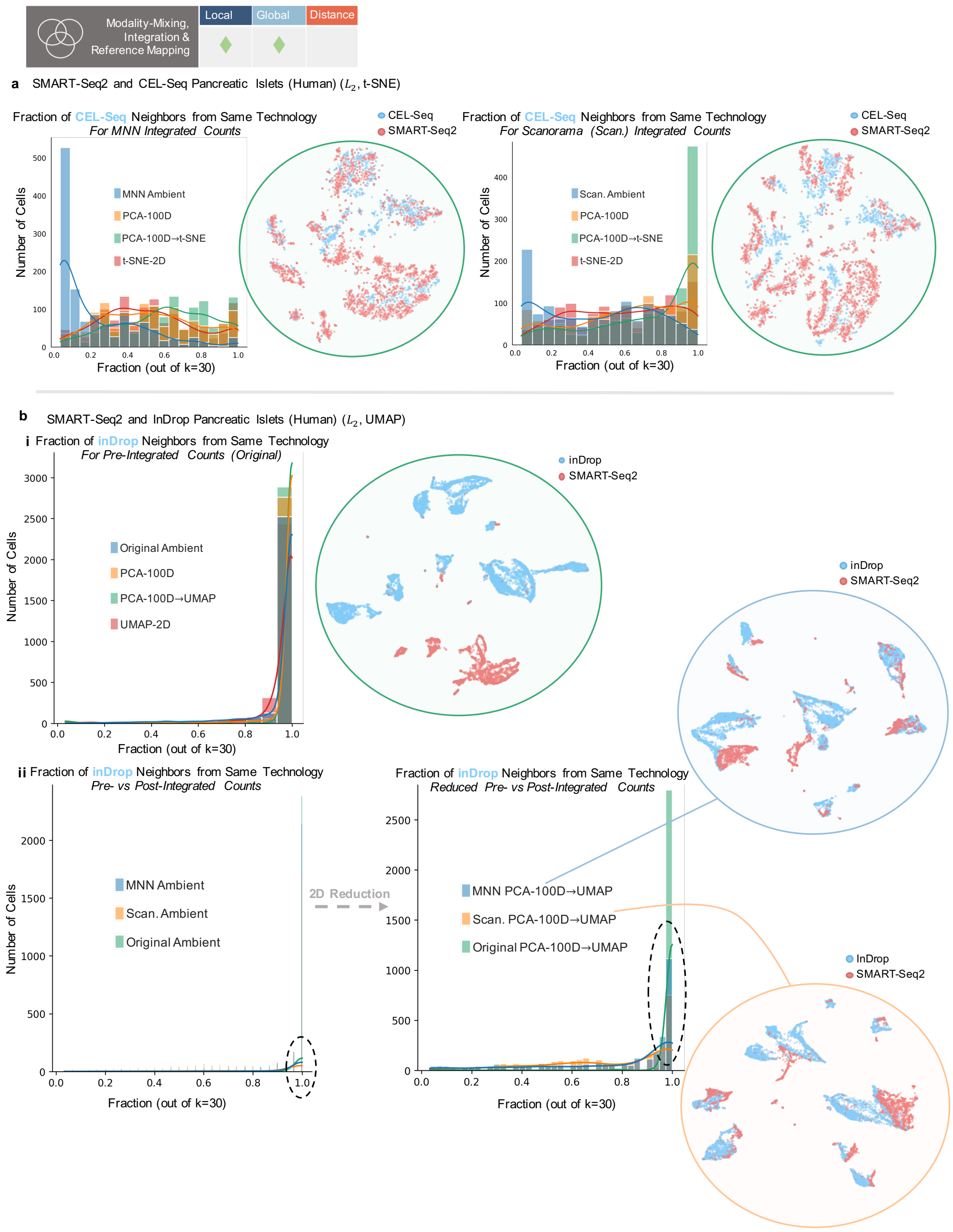

**Supplementary Figure 7: Extended Mixing Pattern Analysis. a)** Distributions of mixing fractions of cells in latent spaces, for integrated CEL-Seq cells, using MNN (left) or Scanorama (right). t-SNE use to make 2D embeddings. **b)** i. For SMART-Seq2 and inDrop pancreatic islet cells pre-integration, fraction of mixing shown for ambient and embedding spaces. ii. Distributions of mixing fractions for ambient integration spaces (MNN or Scanorama) and pre-integrated space (Original). (Right) Distributions of mixing fractions for 2D embeddings of MNN-integrated, Scanorama-integrated, and non-integrated data.

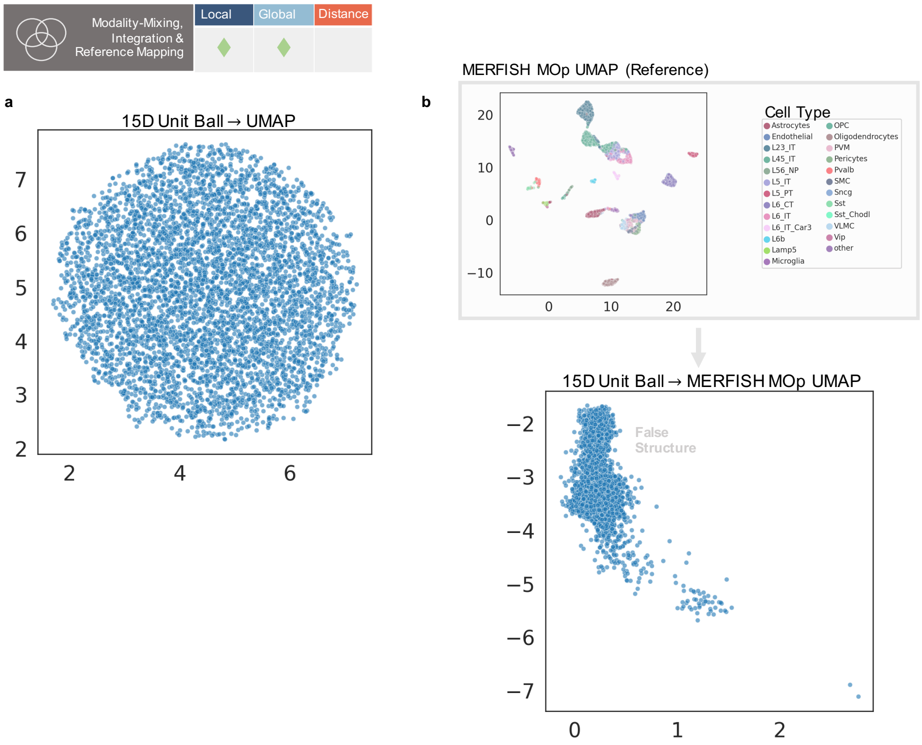

**Supplementary Figure 8: False Imposition of Structure by UMAP Reference Mapping.** **a)** Default UMAP reduction to 2D of 5000 15-dimensional uniformly distributed points (around the unit ball). **b)** (Top) The 2D UMAP on the 15D PCA of the MERFISH MOp dataset. (Bottom) The UMAP transform of the (unseen) 15D unit ball points using the MERFISH UMAP coordinates.

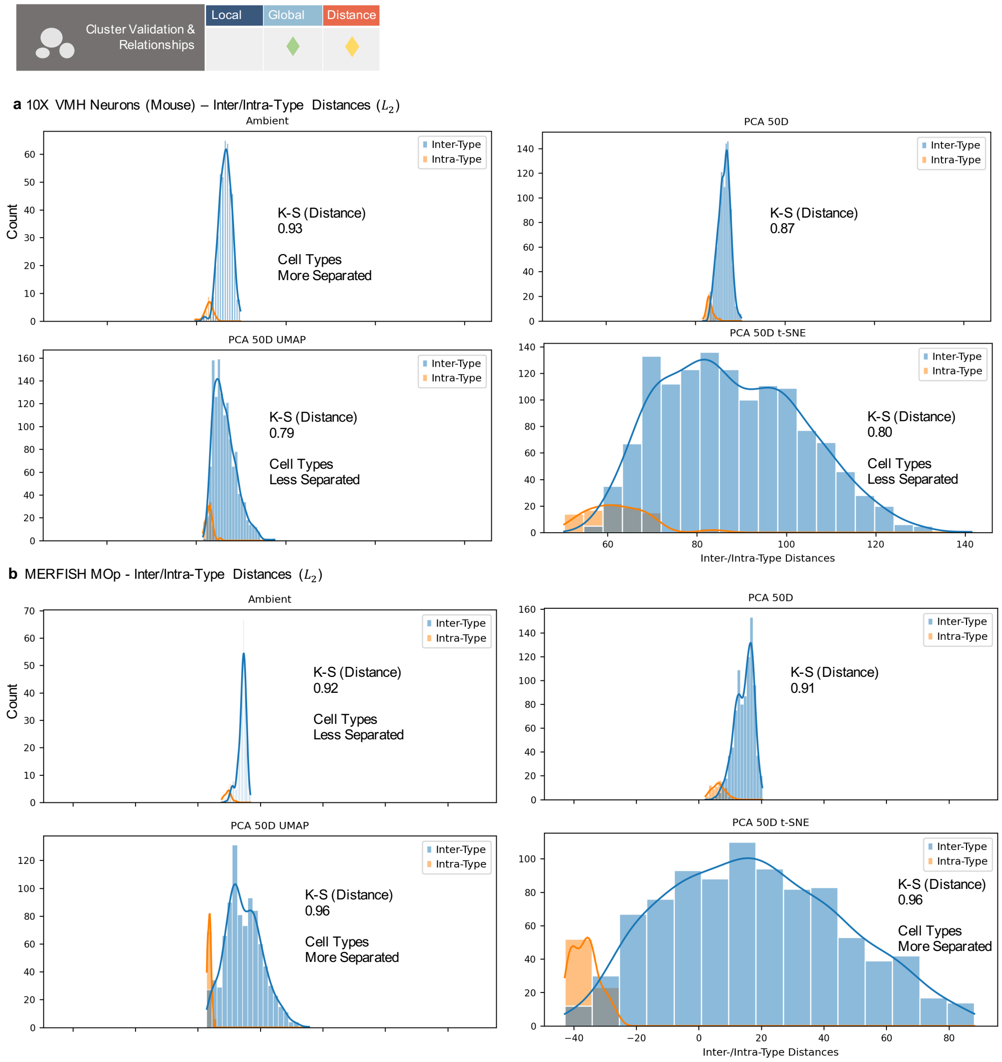

**Supplementary Figure 9: Effect of Reduction on Neighbor Rankings with $L_2$. a)** Two-sample Kolmogorov–Smirnov test statistic for measuring distance/separation between the two distributions of all pairwise inter- or intra-type distances in the 10x VMH data. Distributions shown for all intermediate and non-coupled embeddings (scaled to the same mean for comparison). **b)** Two-sample Kolmogorov–Smirnov test statistic for measuring distance/separation between the two distributions of all pairwise inter- or intra-type distances in the MERFISH MOp data. Distributions shown for all intermediate and non-coupled embeddings (scaled to the same mean for comparison). All calculations use $L_2$ distance.

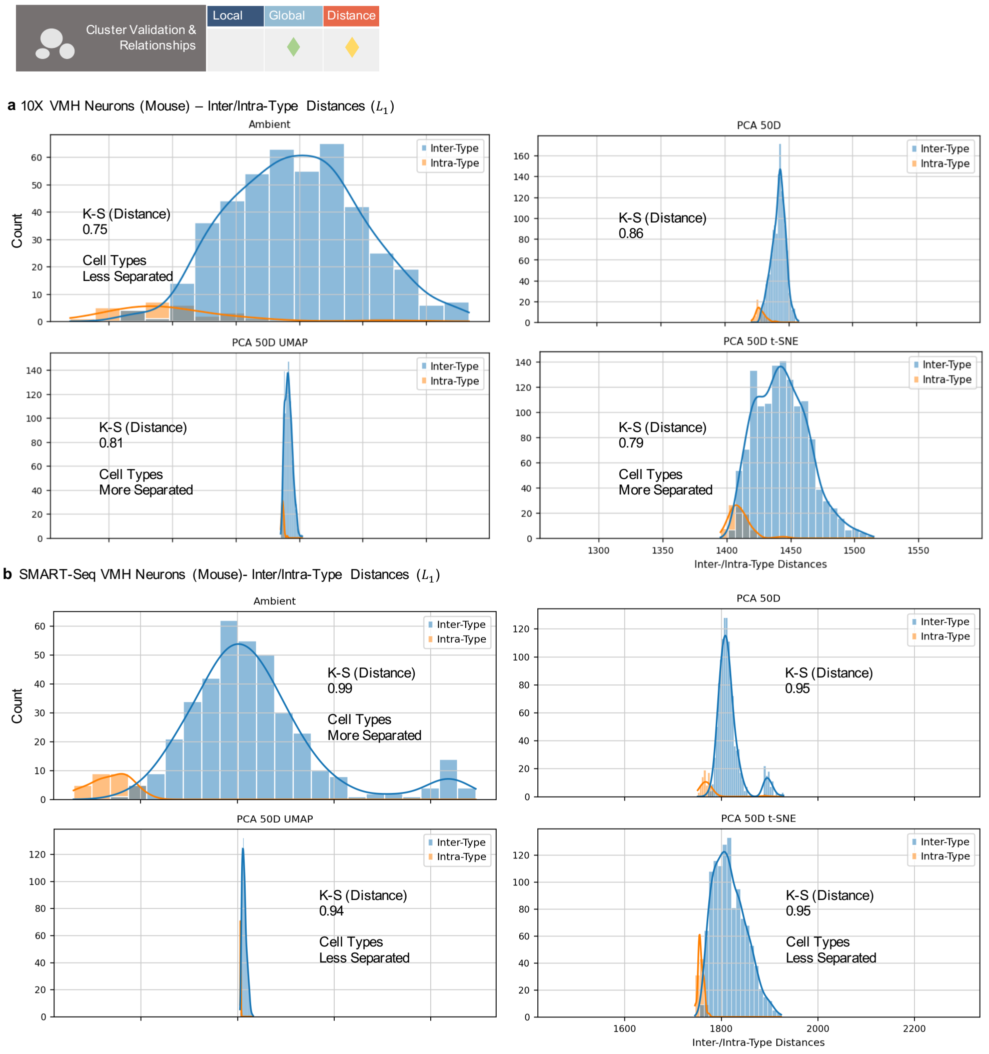

**Supplementary Figure 10: Effect of Reduction on Neighbor Rankings with $L_1$. a)** Two-sample Kolmogorov–Smirnov test statistic for measuring distance/separation between the two distributions of all pairwise inter- or intra-type distances in the 10x VMH data. Distributions shown for all intermediate and non-coupled embeddings (scaled to the same mean for comparison). **b)** Two-sample Kolmogorov–Smirnov test statistic for measuring distance/separation between the two distributions of all pairwise inter- or intra-type distances in the SMART-Seq VMH data. Distributions shown for all intermediate and non-coupled embeddings (scaled to the same mean for comparison). All calculations use $L_1$ distance.

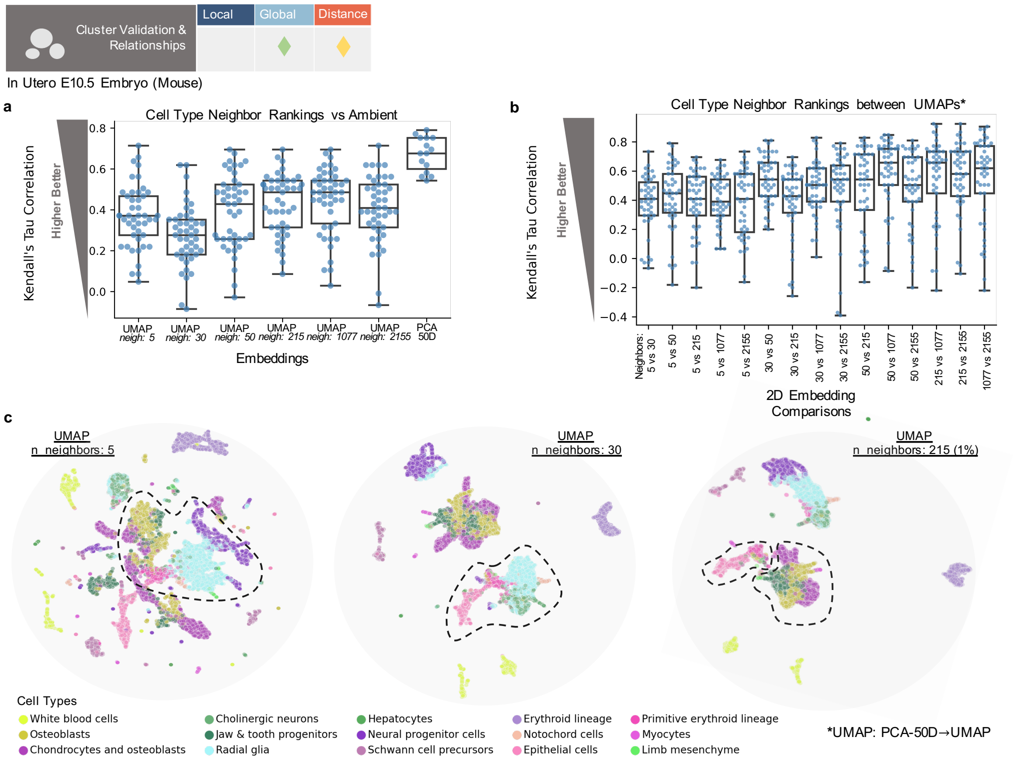

**Supplementary Figure 11: Malleability of Cluster Relationships. a)** Kendall's Tau correlation of cell type neighbor rankings (of in-utero E10.5 embedded data) to ambient data. Whiskers denote 1.5 times the IQR. Plots for n=3 different rounds of UMAP embeddings. **f)** Kendall's Tau correlation of cell type neighbor rankings (of in-utero E10.5 embedded data) between UMAP embeddings. Whiskers denote 1.5 times the IQR. Plots for n=3 different rounds of UMAP embeddings. **g)** PCA-50D→UMAP embedding of in-utero data, with increasing n_neighbors (UMAP parameter) (from left to right). Example of differing cell-cell/cluster relations enclosed in black dashed circles.

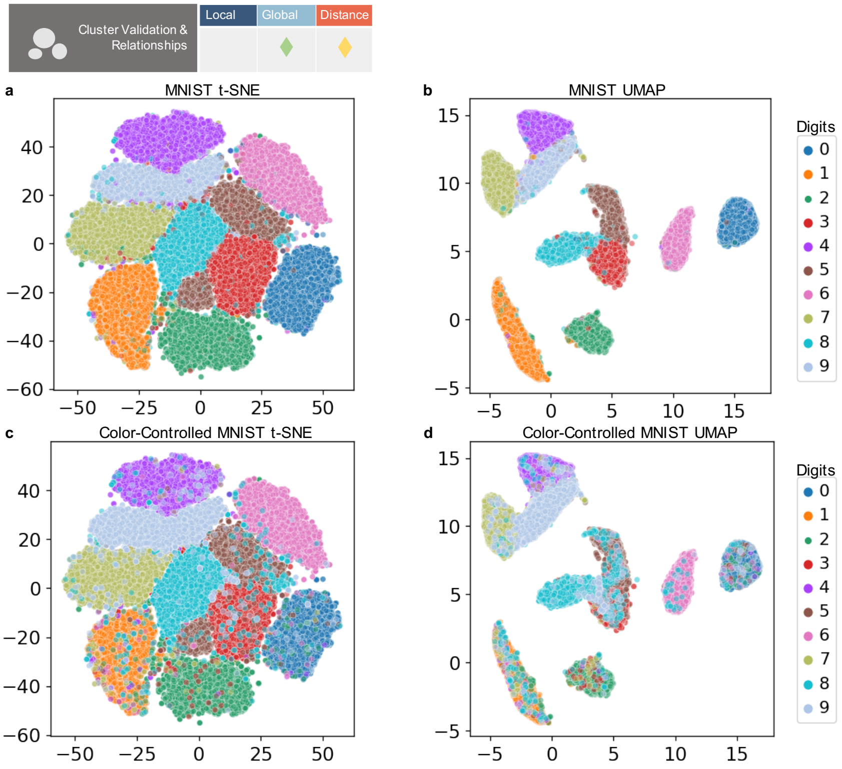

**e** Fraction Correct Digits in t-SNE Clusters

| Digit | Fraction |
|---|---|
| 0 | 0.991221 |
| 1 | 0.910264 |
| 2 | 0.945787 |
| 3 | 0.822215 |
| 4 | 0.611092 |
| 5 | 0.678104 |
| 6 | 0.983947 |
| 7 | 0.833839 |
| 8 | 0.829944 |
| 9 | 0.569507 |

**f** Fraction Correct Digits in UMAP Clusters

| Digit | Fraction |
|---|---|
| 0 | 0.992234 |
| 1 | 0.549540 |
| 2 | 0.955690 |
| 3 | 0.967705 |
| 4 | 0.981684 |
| 5 | 0.919019 |
| 6 | 0.989861 |
| 7 | 0.973025 |
| 8 | 0.873526 |
| 9 | 0.696756 |

**g** Correlation to Ambient Intra-Distances

| Latent | Pearsonr |
|---|---|
| PCA 50D | 0.994942 |
| t-SNE | 0.103368 |
| UMAP | -0.750536 |
| PCA 2D | 0.882780 |

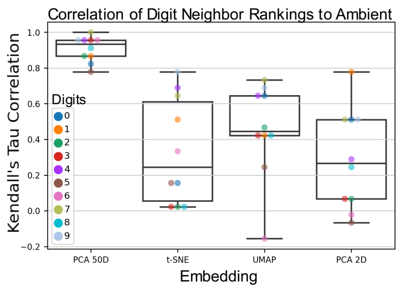

**Supplementary Figure 12: MNIST Embedding Properties. a)** Default t-SNE of the MNIST dataset. **b)** Default UMAP of the MNIST dataset. **c)** t-SNE MNIST plot with hidden points plotted in reverse order. **d)** UMAP MNIST plot with hidden points plotted in reverse order. **e)** Fraction of the correct digit in each of the ten k-means clusters from the t-SNE embedding (see Supplementary Methods). **f)** Fraction of the correct digit in each of the ten k-means clusters from the UMAP embedding. **g)** Pearsonr correlation of intra-distances (internal variance) of each digit, in each embedding, to the ambient variances. **h)** Kendall's Tau correlation of each digit's neighbor rankings to ambient space. For box plots, whiskers denote 1.5 times the IQR for the displayed embedding. The 50D PCA space was not coupled to the 2D embedding/reductions, as this is not standard practice for the MNIST dataset.

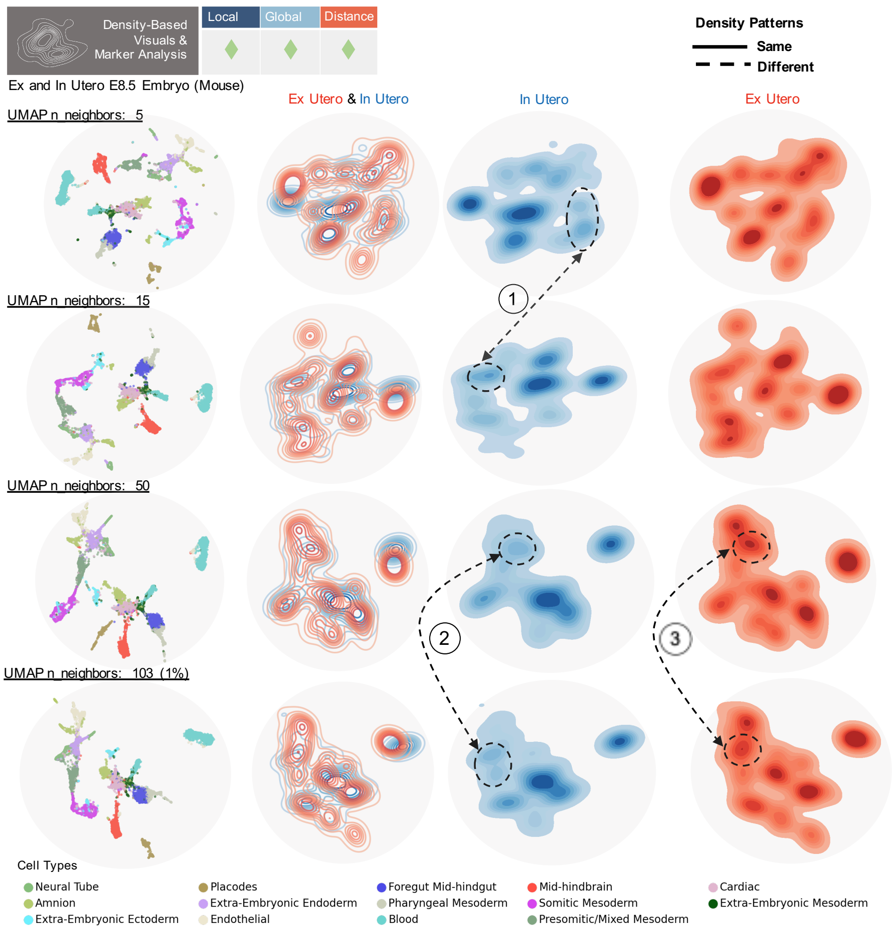

**Supplementary Figure 13: Extended Analysis of Density-based Visuals for UMAP a)** Top row (left to right) displays UMAP embedding with 5 neighbors, embedding contour plot colored by condition, embedding of just in-utero cells, embedding of just ex-utero cells. UMAP n_neighbors increases for each row (5,15,50, and 103 neighbors down the rows). Numbers denote comparisons between plots, dashed lines denote a difference, and solid lines denote the same appearance.

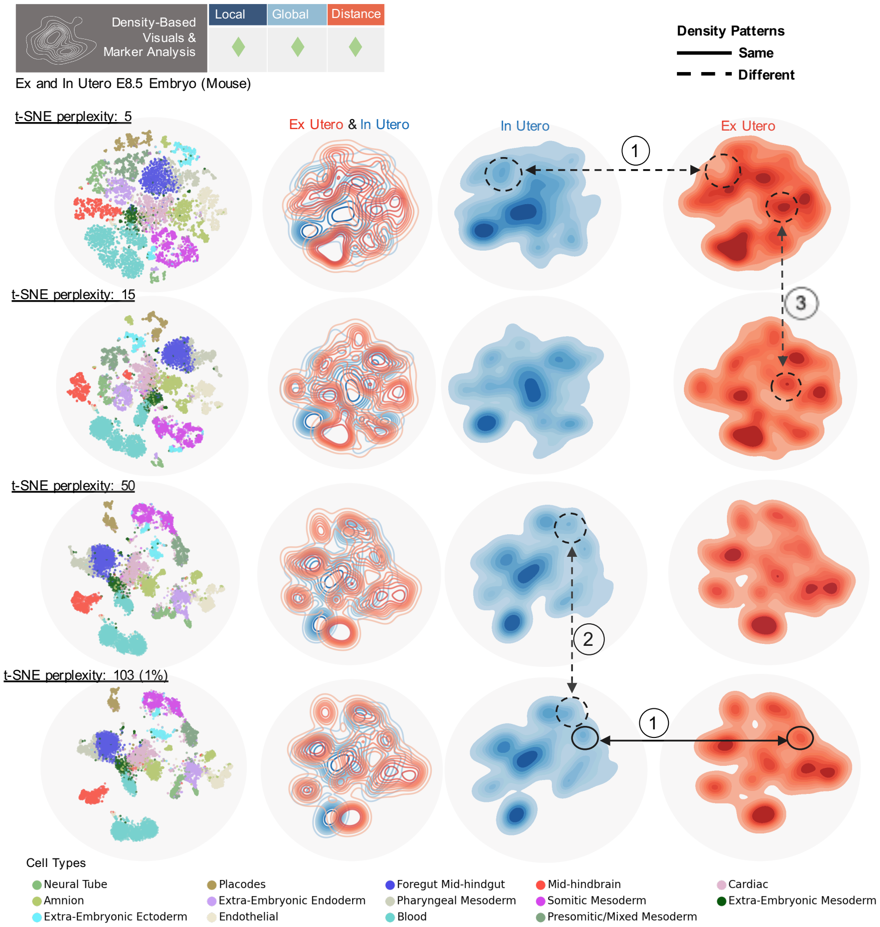

**Supplementary Figure 14: Extended Analysis of Density-based Visuals for t-SNE a)** Top row (left to right) displays t-SNE embedding with 5 neighbors, embedding contour plot colored by condition, embedding of just in-utero cells, embedding of just ex-utero cells. T-SNE perplexityincreases for each row (5, 15, 50, and 103 neighbors down the rows). Numbers denote comparisons between plots, dashed lines denote a difference, and solid lines denote the same appearance.

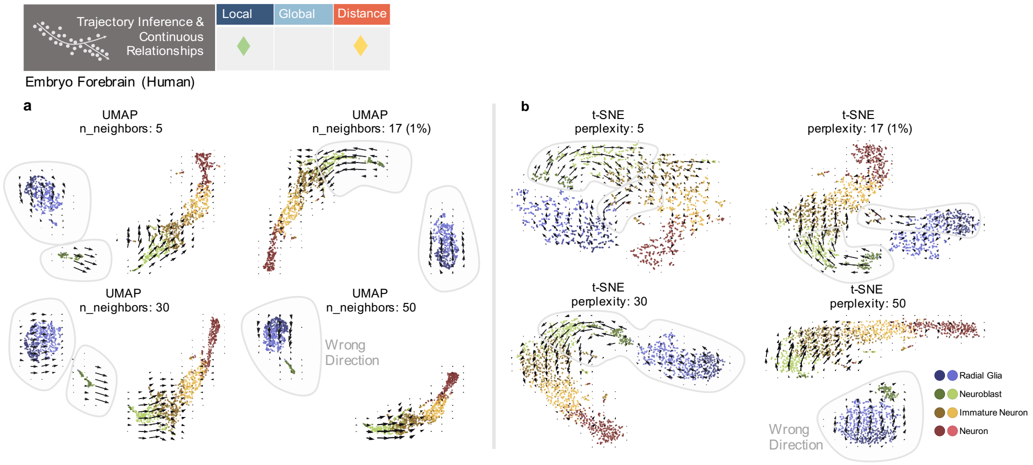

**Supplementary Figure 15: Extended Analysis of Trajectory Inference with 2D Embeddings a)** Velocyto RNA velocity embeddings for UMAPs made with 5,17, 30 or 50 n_neighbors. Cell types of interest highlighted in grey. **b)** Velocyto RNA velocity embeddings for t-SNEs made with 5, 17, 30 or 50 perplexity values. Cell types of interest highlighted in grey.

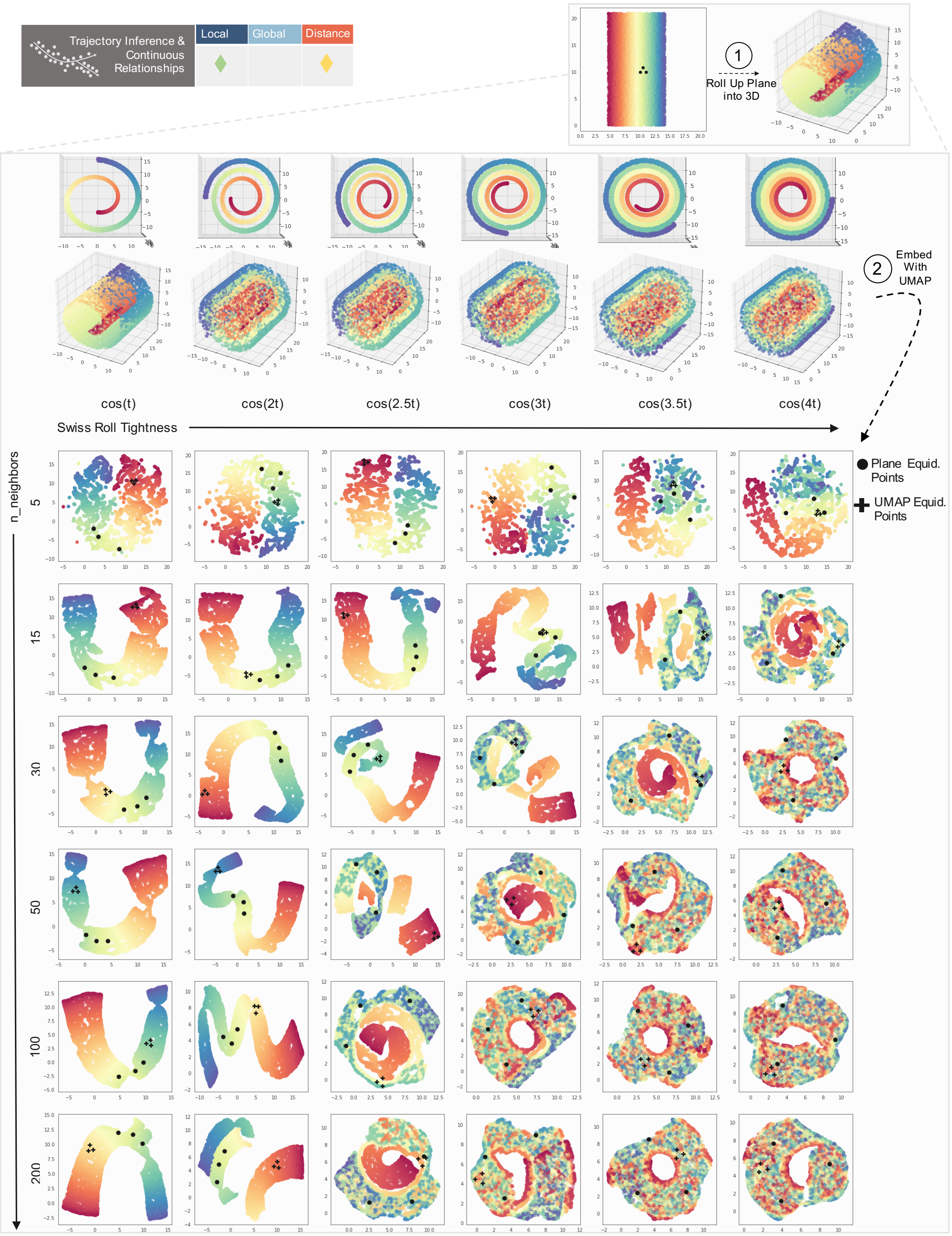

**Supplementary Figure 16: Swiss Roll UMAP Embeddings over Parameter Grid.** (1) denotes how from randomly generated 2D plane of 10,000 points, with three additional equidistant points, a 3D Swiss Roll is constructed. Black dots denote the original three equidistant points. See Supplementary Methods. (2) Across the x direction is tightness of the roll, and down the y direction is the n_neighbors parameter used for UMAP embedding. Topmost row of spirals shows head-on view of same 3D spirals in row below. The UMAPs that comprise the grid are the 2D UMAP embeddings of the corresponding 3D Swiss roll with n_neighbors parameter.

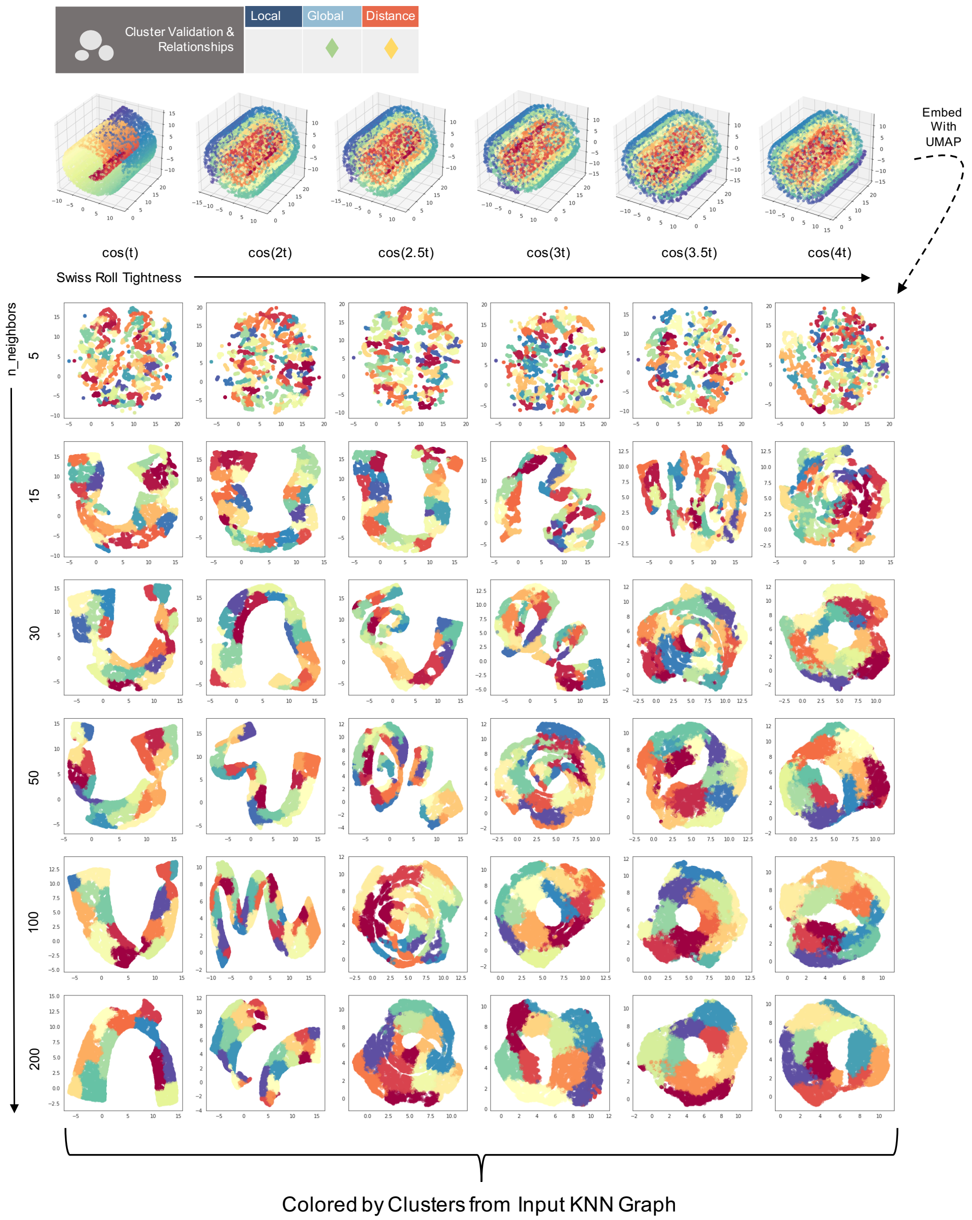

**Supplementary Figure 17: Louvain Clustering of Swiss Roll UMAPs.** Default Louvain clustering using the same neighborhood graph as input into the UMAP algorithm, colored by assigned Louvain clusters (not original/2D plane color assignments).

#### Picasso & MCML Results

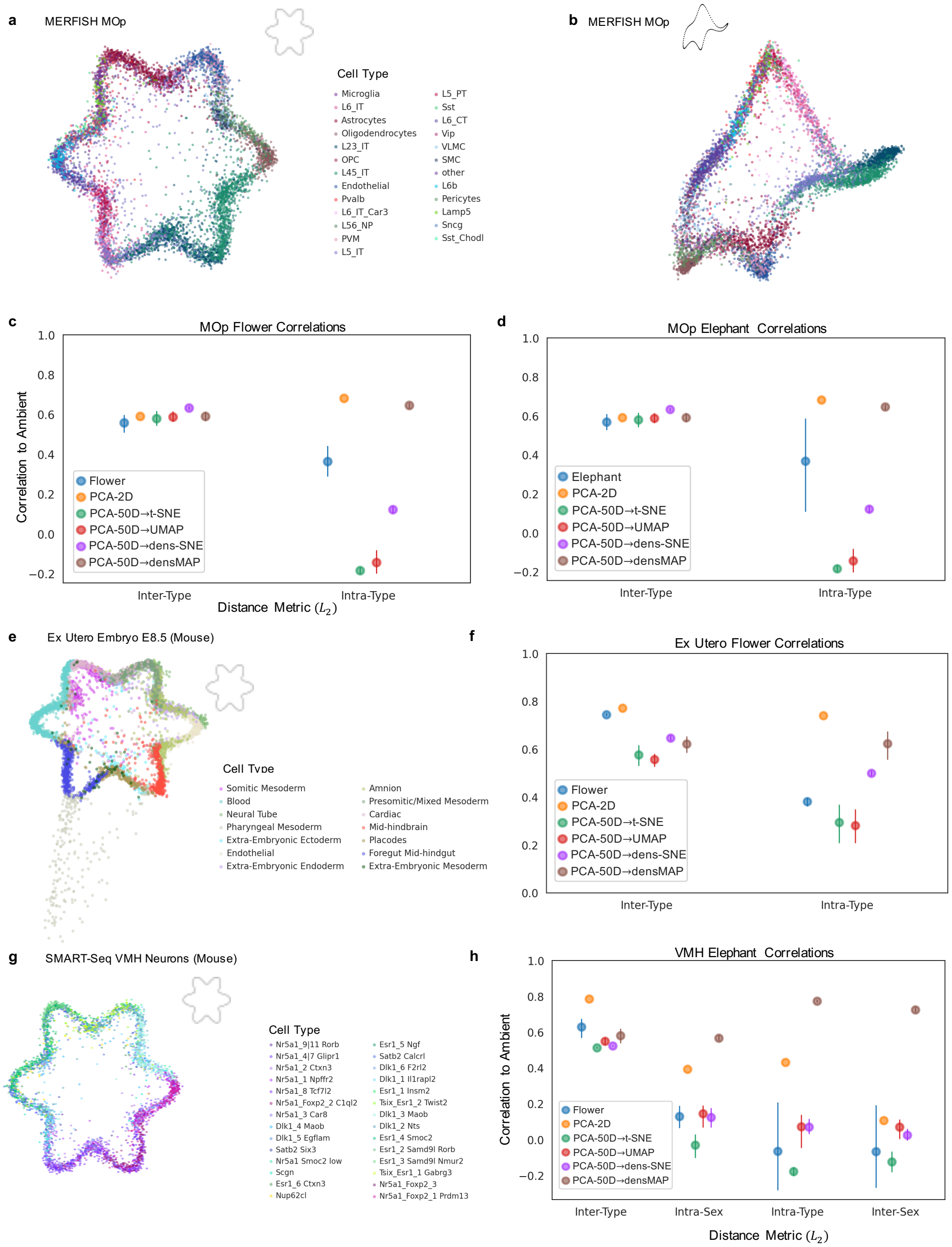

**Supplementary Figure 18: Further Analysis of Picasso Embedding with $L_2$. a)** Picasso embedding of the MERFISH MOp data fit to a flower-like boundary. **b)** Picasso embedding of the MOp data fit to a 'von Neumann' elephant. **c)** Comparison of correlation metrics between the flower Picasso embedding and the other baseline 2D embeddings, including densVis [61] embeddings. **d)** Comparison of correlation metrics between the elephant Picasso embedding and the other baseline 2D embeddings, including densVis embeddings. **e)** Picasso embedding of the ex-utero mouse Embryo E8.5 data fit to a flower-like boundary. **f)** Comparison of correlation metrics between the flower Picasso embedding and the other baseline 2D embeddings, including densVis embeddings. **g)** Picasso embedding of the SMART-Seq mouse VMH neurons dataset fit to a flower-like boundary. **h)** Comparison of correlation metrics between the flower Picasso embedding and the other baseline 2D embeddings, including densVis embeddings. For all plots bars denote the 95% C.I. and were run over 5 rounds of generated embeddings. All calculations done with $L_2$ distance.

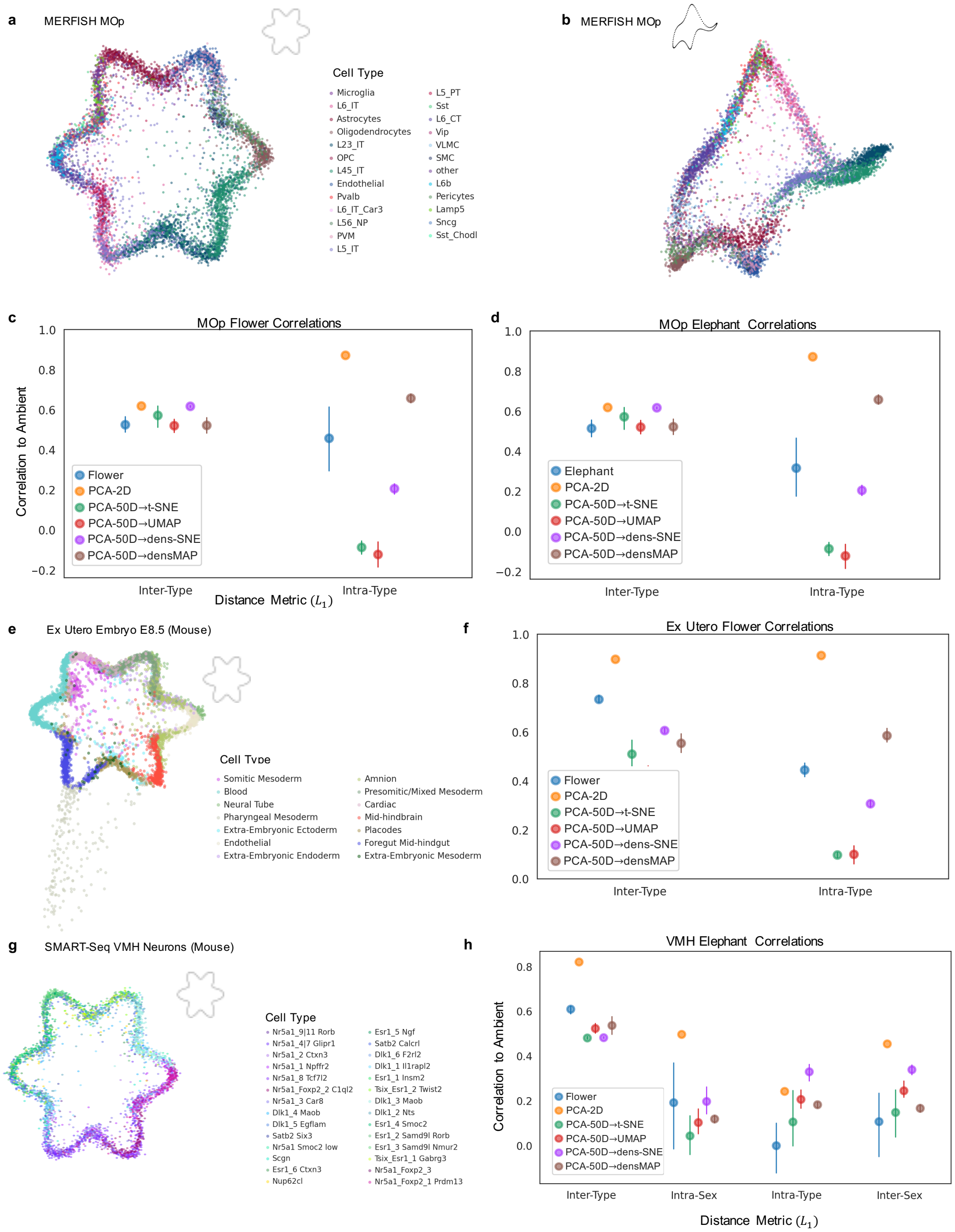

**Supplementary Figure 19: Further Analysis of Picasso Embedding with $L_1$. a)** Picasso embedding of the MERFISH MOp data fit to a flower-like boundary. **b)** Picasso embedding of the MOp data fit to a 'von Neumann' elephant. **c)** Comparison of correlation metrics between the flower Picasso embedding and the other baseline 2D embeddings, including densVis embeddings. **d)** Comparison of correlation metrics between the elephant Picasso embedding and the other baseline 2D embeddings, including densVis embeddings. **e)** Picasso embedding of the ex-utero mouse Embryo E8.5 data fit to a flower-like boundary. **f)** Comparison of correlation metrics between the flower Picasso embedding and the other baseline 2D embeddings, including densVis embeddings. **g)** Picasso embedding of the SMART-Seq mouse VMH neurons dataset fit to a flower-like boundary. **h)** Comparison of correlation metrics between the flower Picasso embedding and the other baseline 2D embeddings, including densVis embeddings. For all plots bars denote the 95% C.I. and were run over 5 rounds of generated embeddings. All calculations done with $L_1$ distance.

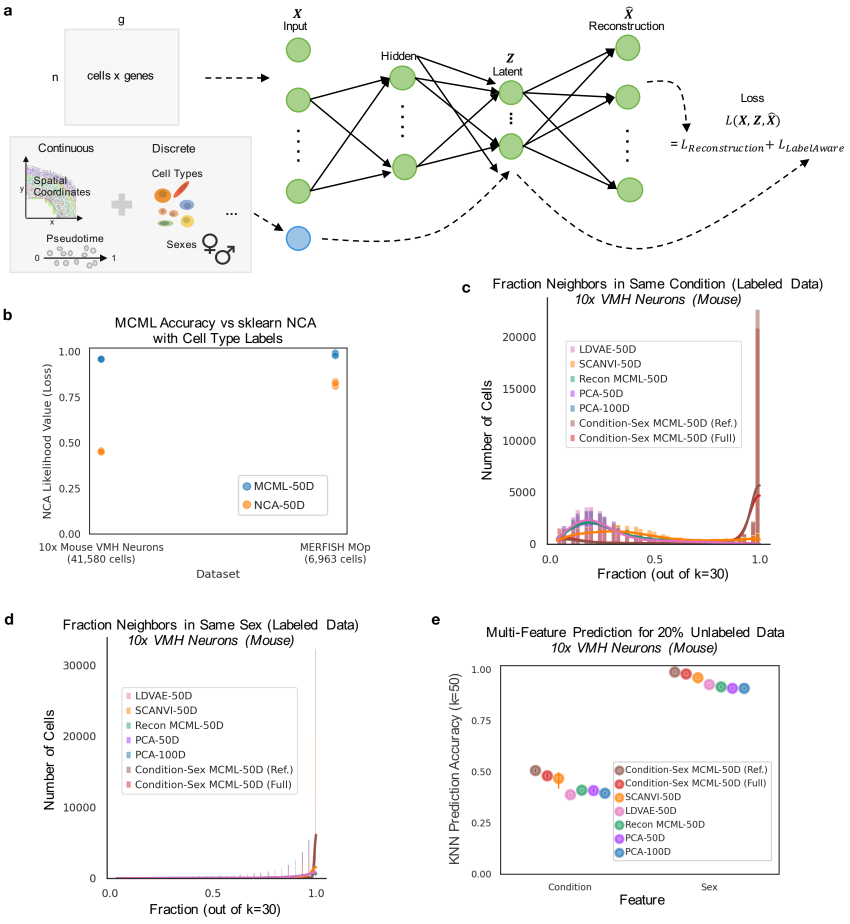

**Supplementary Figure 20: Multi-Class, Multi-Label Embedding (MCML). a)** Diagram of input data with various labels, autoencoder structure, and loss function, to create embedding $Z$. **b)** Comparison of MCML with no reconstruction (equivalent to the NCA algorithm) with sklearn NCA implementation with cell type labels. Measured by NCA likelihood objective (higher better) (see Supplementary Methods). **c)** Fraction of neighbors with the same label across embeddings. Measured for behavioral condition labels. **d)** Fraction of neighbors with the same label across embeddings. Measured for sex labels. **e)** KNN prediction accuracy for embeddings, for predicting condition or sex label of unlabeled cells.

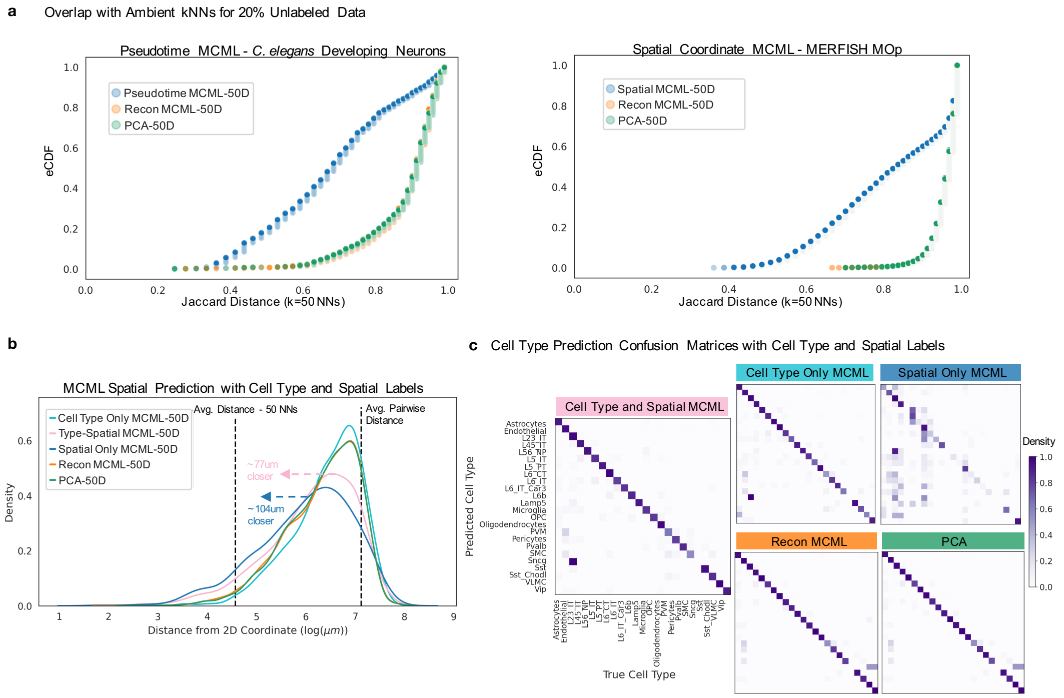

**Supplementary Figure 21: Embedding Continuous and Discrete Labels. a)** (Left) Jaccard distance between neighbors in embedded spaces and pseudotime neighbors from ambient space. (Right) Jaccard distance between neighbors in embedded spaces and true spatial neighbors. **b)** Distance of spatial location KNN predictions (for unlabeled cells) from true locations. Average prediction improvements denoted for Spatial and Type-Spatial MCML embeddings, over other denoted embeddings. **c)** Confusion matrices depicting clarity of cell type classification using NNs in each embedding space (see Supplementary Methods).

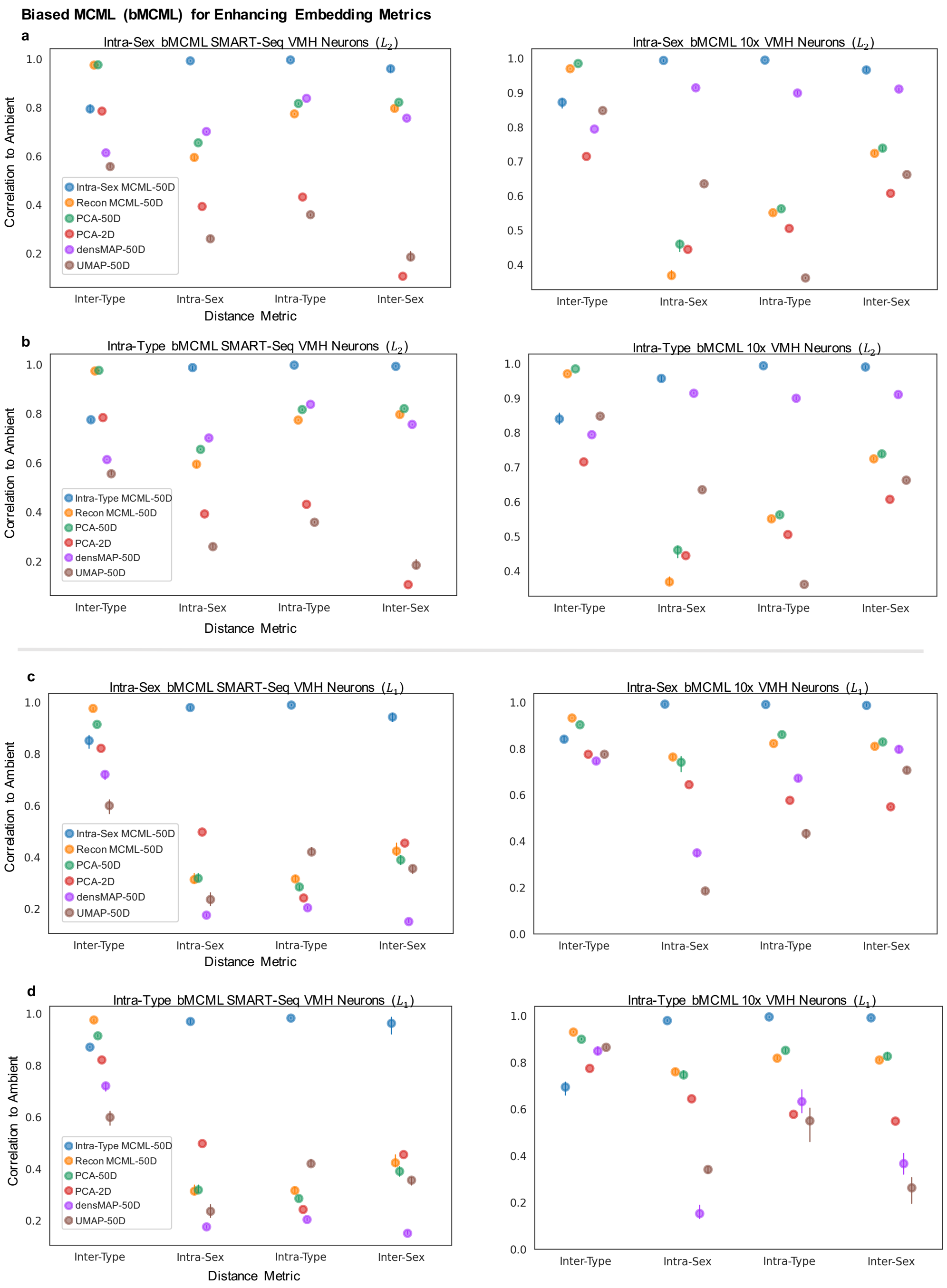

**Supplementary Figure 22: Recapitulation of Ambient Properties with bMCML. a)** Correlation metrics for bMCML (Biased MCML) with intra-sex distance correlation as the objective function, for the SMART-Seq mouse VMH neurons (left) and the 10x VMH Neurons (right). (See Supplementary Methods). **b)** Correlation metrics for bMCML (Biased MCML) with intra-type distance correlation as the objective function, for the SMART-Seq mouse VMH neurons (left) and the 10x VMH Neurons (right). **c)** Same correlation plots as **a**, but using $L_1$ distance instead of $L_2$. **d)** Same correlation plots as **b**, but using $L_1$ distance instead of $L_2$. For all plots bars denote the 95% C.I.

#### Supplementary Note

##### Bounds on Distortion of Equidistant Points

Induced distortion has been investigated in the literature for various conformations and embedding of points, e.g. the minimum distortion bound for embedding an $n$-point spherical metric onto a line [40] (akin to pseudotime inference), and the number of dimensions required to embed a metric space into a low-dimension normed space (defined by some $l$-norm) [41]. However, investigation of the implication of these bounds in real datasets across the sciences has been limited. Here we focus on the case of equidistant points and their distortion in two-dimensions to provide a more concrete realization of such bounds in the context of single-cell gene expression.

A trivial case is the result that no more than three points can be equidistant points in $\mathbb{R}^2$ (no more than $n+1$ points in $\mathbb{R}^n$). This raises the question of how close to equidistant more than three points in $\mathbb{R}^2$ can be as even near-equality is impossible; specifically, a lower bound on the ratio between the maximum and minimum pairwise distances shows that distortion, which increases with the number of points, is inevitable.

A straightforward way to see this is via the two-dimensional isodiametric inequality which states that among all shapes of a given diameter, the circle has the greatest area (for a simple proof see [46]). Formally, for any body in $\mathbb{R}^2$, the area $A$ is bounded above by $\frac{\pi}{4}$ times the square of the diameter $D$ (the supremum of distances between any pair of points), i.e.

$$A \leq \frac{\pi}{4} D^2. \tag{4}$$

**Theorem 1** *Given $n \geq 3$ points in $\mathbb{R}^2$, let $d$ be the minimum distance among all pairs of points, and $D$ the maximum distance (i.e. the diameter). The ratio of $D$ to $d$ satisfies*

$$\frac{D}{d} \geq \sqrt{\frac{n-2}{2}}. \tag{5}$$

**Proof**: Let $B$ be the set of points consisting of the convex hull of $n$ points in $\mathbb{R}^2$, and let $I$ denote the remaining points, with $|B| = k$ and $|I| = n - k$. Note that for each point in $I$, there exists a semi-circle of radius $\frac{d}{2}$ centered at the point that does not touch any other point, or extend beyond the convex hull of the points (Supplementary Note Fig. 1). If we denote the sum of the areas of these semi-circles by $A_I$, we obtain

$$\begin{aligned} A_I &= \frac{1}{2}\left(\pi\left(\frac{d}{2}\right)^2\right)(n-k) \\ &= \frac{\pi d^2}{8}(n-k)\ . \end{aligned}$$

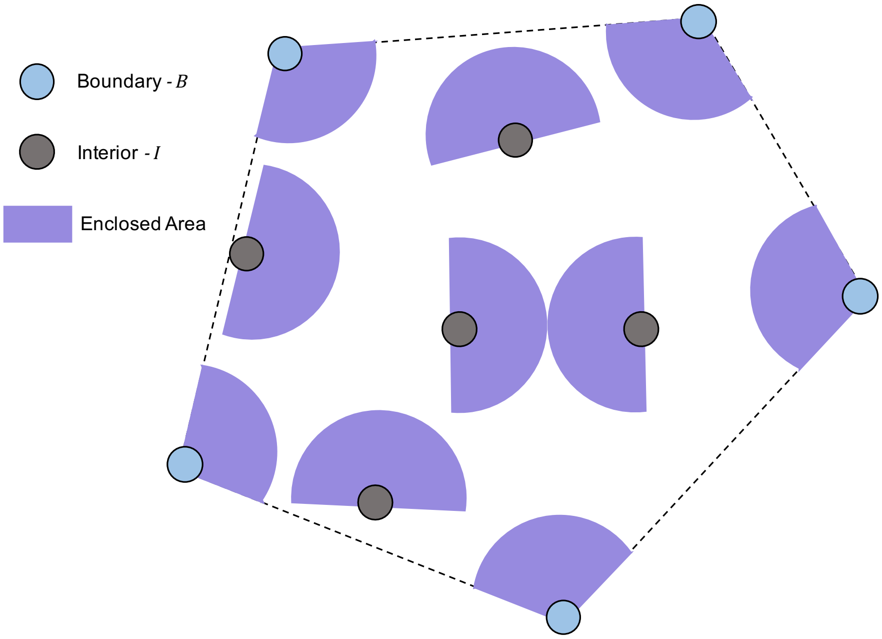

**Supplementary Note Fig. 1: Bounding the Area Enclosed by Points in Two-Dimensions.** Example of a set of 10 points showing the enclosed area for points in the $I$ and $B$ sets in the proof of Theorem 1.

Furthermore, for each of the $k$ points in $B$, there is a circle sector of radius $\frac{d}{2}$ spanning the interior angle of the convex hull at that point that does not touch any other point, or extend beyond the convex hull. Since the sum of the interior angles of a $k$-gon is $(k-2)\pi$, we find that the sum of the areas of the circle sectors, which we denote by $A_B$, is given by

$$\begin{aligned} A_B &= \pi \left(\frac{d}{2}\right)^2 \left(\frac{(k-2)\pi}{2\pi}\right) \\ &= \frac{\pi d^2}{8}(k-2) \ . \end{aligned}$$

Summing $A_I$ and $A_B$, we obtain a bound for the area enclosed by the $n$ points:

$$\begin{aligned} A &\geq A_I + A_B \\ &= \frac{\pi d^2}{8}(k-2) + \frac{\pi d^2}{8}(n-k) \\ &= \frac{\pi d^2}{8}(n-2) \ . \end{aligned} \tag{6}$$

Combining the upper (4) and lower (6) bounds for the area $A$, we find that

$$\begin{aligned} \pi \frac{D^2}{4} &\geq \frac{\pi d^2}{8}(n-2) \\ \Rightarrow \frac{D}{d} &\geq \sqrt{\frac{n-2}{2}} \ . \end{aligned} \tag{7}$$
